# Sense and Screen-ability: Development of tuneable, biosensor-based screening platforms for abscisic acid

**DOI:** 10.1101/2023.05.16.540971

**Authors:** Maximilian Otto, Yasaman Dabirian, Florian David, Verena Siewers

## Abstract

The activities of heterologous enzymes often limit the production titers, rates and yields of cell factories. With the help of biosensors, large random mutagenesis libraries can be screened for improved enzyme variants in a high-throughput manner, even if the enzyme-of-interest is poorly characterised.

We previously constructed a *Saccharomyces cerevisiae* cell factory for the heterologous production of abscisic acid (ABA), a high-value product with a broad range of applications in medicine, agriculture and nutrition. In the current study, we developed high-throughput screening platform strains for two rate-limiting cytochrome P450 monooxygenases, BcABA1 and BcABA2, in the ABA biosynthetic pathway. The screening platforms are designed to minimize the occurrence of false positives during screening experiments.

We thoroughly characterised two plant protein-based ABA biosensor candidates. Furthermore, we designed a simple genetic switch, based on the thiamine-repressible promoter p*THI4*, to regulate the expression level of enzyme variants. We demonstrated that ABA production can be fine-tuned by varying thiamine concentration in the media. In-depth analysis of the platform strains revealed that screening conditions can be optimized by varying thiamine concentration and cultivation time, making it possible to utilize the full dynamic and operational range of the biosensor. In the future, the constructed strains can be used to screen for improved BcABA1 and BcABA2 variants.

## Introduction

The past decade has seen a global effort towards a more sustainable, bio-based economy. Cell factories will play a critical role in this transition by enabling the production of complex biomolecules used in medicine, nutrition, agriculture and cosmetics. Nonetheless, reaching commercially relevant product titers, rates and yields remains a challenge and currently hinders the wide-scale adoption of cell factories. Many high-value products are secondary metabolites that are usually present in low concentrations in the native host and the relevant pathway enzymes generally exhibit low catalytic activities (Bar-Even et al., 2011). Consequently, enzyme activities often limit production of heterologous metabolites and are one of the central engineering targets in biotechnology (Katsimpouras and Stephanopoulos, 2021).

In a previous study, we developed a proof-of-concept abscisic acid (ABA) cell factory using the biotechnological workhorse *Saccharomyces cerevisiae* (Otto et al., 2019). The sesquiterpenoid (C_15_ terpenoid) ABA has a multitude of applications in agriculture, nutrition and medicine. Historically known for being an essential phytohormone, ABA regulates various developmental processes and is central to the abiotic stress response in higher plants (Ruth Finkelstein, 2013). More recently, the role of ABA in humans has been investigated. Notably, ABA could be used in treatments for inflammatory diseases, pathogen infections, type-2-diabetes and metabolic syndrome (Kim et al., 2020; Lievens et al., 2017; Sakthivel et al., 2016). Potential applications as a bitter taste receptor blocker are also being investigated (Kim et al., 2020; Pydi et al., 2015; Singh et al., 2022).

The yeast cell factory expresses four enzymes of the ABA biosynthetic pathway from the plant pathogenic fungus *Botrytis cinerea* (Otto et al., 2019; Siewers et al., 2006, 2004; Takino et al., 2019). The native precursor farnesyl pyrophosphate (FPP) is first cyclised by BcABA3, and the ABA core scaffold is thereafter oxidized at multiple positions by two cytochrome P450 monooxygenases (CYPs), BcABA1 and BcABA2, and the short-chain dehydrogenase/reductase BcABA4 (Figure 1A). We demonstrated that the two CYPs BcABA1 and BcABA2, are limiting ABA production at this stage (Otto et al., 2019). By engineering the CYPs and improving their activity, this production bottleneck could be alleviated.

**Figure 1:**
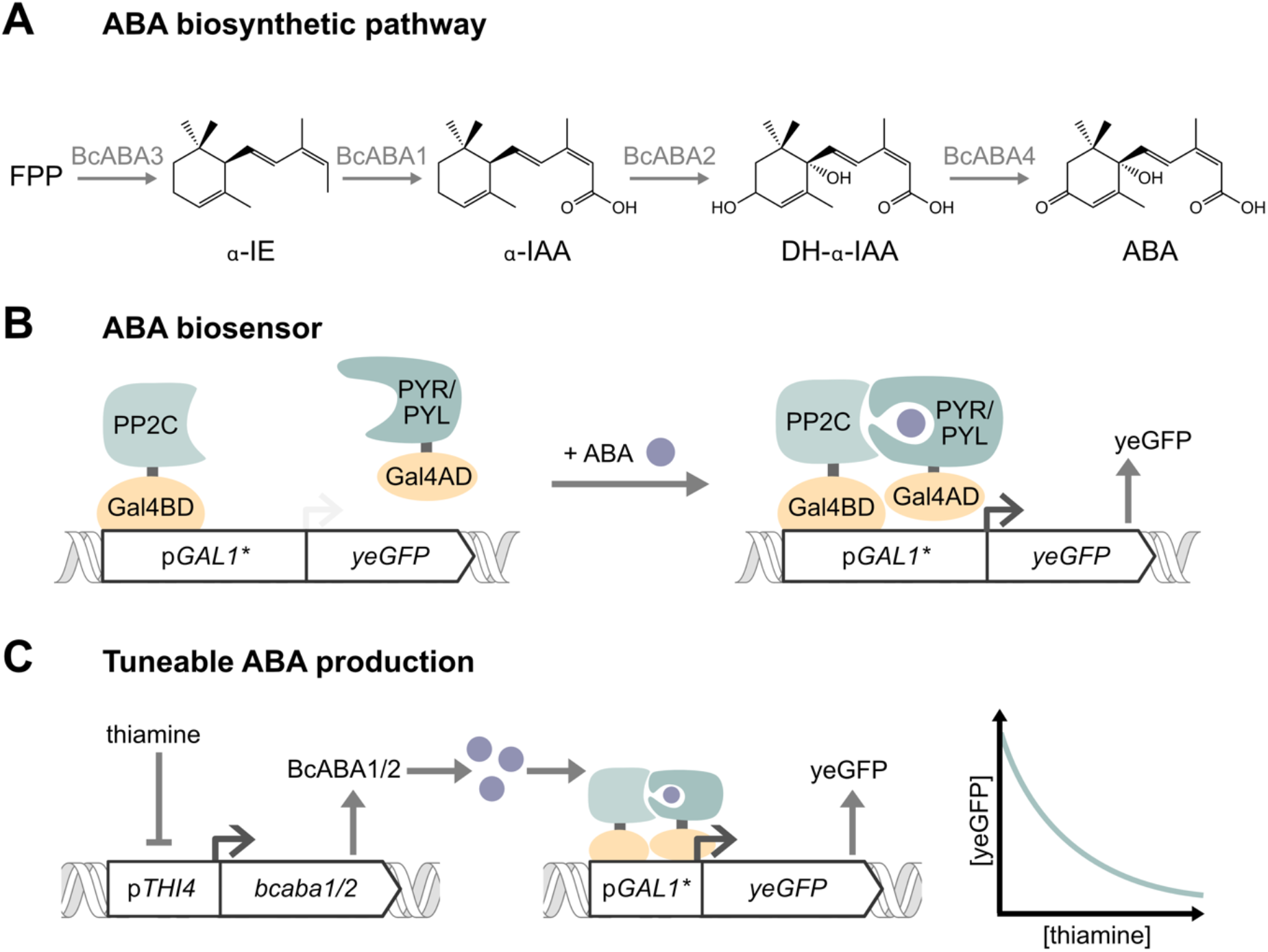
Project overview highlighting different genetic modules of the screening platform strains. **A**: ABA biosynthetic pathway from *B. cinerea* introduced in *S. cerevisiae*. The pathway consists of the four genes *bcaba1*, *bcaba2*, *bcaba3* and *bcaba4*. In addition to the pathway genes, the *B. cinerea* cytochrome P450 reductase gene, *bccpr1*, is expressed since *bcaba1* and *bcaba2* encode CYPs. FPP is the native, universal sesquiterpene precursor. **B**: Schematic of the Y2H-based ABA biosensor. Green shapes represent *A. thaliana* proteins and orange shapes represent *S. cerevisiae* proteins. The biosensor consists of two fusion proteins. One fusion protein contains the Gal4BD and a PP2C variant from *A. thaliana*. The other fusion protein contains the Gal4AD and a variant of the PYR/PYL ABA receptor family from *A. thaliana.* The reporter cassette consists of a modified *GAL1* promoter in which the Mig1 binding sites were deleted and *yeGFP* as the reporter gene. In the presence of ABA, the two fusion proteins co-localize at the reporter cassette p*GAL1\**-*yeGFP* and the fluorescent reporter is being expressed. **C**: Schematic of *pTHI4*-mediated ABA production and biosensor output in the screening platform strains. *Bcaba1* or *bcaba2* are regulated by the thiamine-repressible promoter p*THI4*. In the absence of thiamine or in low concentrations, *bcaba1* or *bcaba2* is highly expressed, leading to high ABA production and high biosensor reporter output. By varying the thiamine concentration, ABA production can be fine-tuned to the operational range of the biosensor and the activity of the enzyme. This tuneable production allows the screening conditions to be optimized. Abbreviations: FPP = farnesyl pyrophosphate, α-IE = α-ionylideneethane, α-IAA = α-ionylideneacetic acid, DH-α-IE = 1ʹ,4ʹ-trans-dihydroxy-α-ionylideneacetic acid, ABA = abscisic acid, Y2H = yeast two-hybrid, BD = binding domain, PP2C: protein phosphatase 2C, yeGFP = yeast enhanced GFP

CYPs are a superfamily of enzymes that can catalyse various oxidation reactions using molecular oxygen (Werck-Reichhart and Feyereisen, 2000). CYPs have been in the focus of research for decades. This is in part due to their important roles in drug and toxin metabolism in humans (Guengerich, 2006), and in addition due to their abundance in biosynthetic pathways in prokaryotic and eukaryotic organisms (Urlacher and Girhard, 2019). CYPs are complex enzymes and often exhibit low catalytic efficiencies (Jung et al., 2011). They contain heme as a co-factor and most eukaryotic CYPs require cytochrome P450 reductases (CPRs) as co-enzymes that provide electrons via NAD(P)H (Werck-Reichhart and Feyereisen, 2000). Eukaryotic CYPs are predominantly membrane-anchored (Werck-Reichhart and Feyereisen, 2000). With regards to the different functional domains and potential co-dependencies between them, CYP engineering remains challenging despite the continued research efforts. Random mutagenesis coupled with high throughput screening (HTS) or selection is a tried-and-tested approach to find improved enzyme variants.

High-throughput screening of mutagenesis libraries requires a detectable signal that correlates with the activity of the enzyme, usually the product concentration. Whole-cell biosensors can be used to quantitatively detect molecules *in vivo* and have become an indispensable tool for HTSs (Kaczmarek and Prather, 2021). Biosensors can translate the concentration of the molecule-of-interest (the ligand of the biosensor, in this case the product) into an easy-to-detect reporter signal, e.g. fluorescence. Subsequently, individual mutants can be discriminated according to their reporter signal and cells with high reporter signal containing improved enzyme variants can be isolated. Fluorescence-activated cell sorting (FACS) is a commonly used technique to screen and sort mutant libraries according to their fluorescent signal.

The requirements for a biosensor are highly application-dependent and biosensor characteristics often require fine-tuning to increase the chances of success (Lim et al., 2018), in this case the chance to find an improved enzyme variant. Two important biosensor characteristics are the dynamic range and the operational range. The dynamic range is the difference between the minimal reporter signal (in the “OFF state”) and the maximal reporter signal (in the “ON state”). The operational range is the concentration range, in which the reporter signal changes. Together, the dynamic and operational range constitute a biosensor’s response curve, the correlation between ligand concentration and reporter signal. Using a biosensor with an unsuitable response curve will result in a large number of false positive or false negative events, potentially making the screening impossible. Changing a biosensoŕs response curve is often not a trivial task and might require iterative rounds of engineering since the dynamic and operational range are interdependent.

In plants, ABA sensing is mediated by the PYR/PYL receptor family (also referred to as RCAR receptors) (Fidler et al., 2022). Upon binding of ABA to a PYR/PYL receptor, a protein interaction domain is exposed, and the receptor forms a heterodimer with a protein phosphatase 2C (PP2C), resulting in the inhibition of the phosphatase. This inhibition triggers a signalling cascade, ultimately leading to a transcriptional response (Ng et al., 2014). Due to the central role of ABA in plants, its signalling is highly sophisticated with a multitude of PYR/PYL receptor and PP2Cs involved. Researchers interested in the ABA signalling pathway first identified the ABA-induced dimerization of PYR/PYLs and PP2Cs in a yeast-2-hybrid (Y2H) assay, in which a PP2C or a PYR/PYL receptor were used as baits (Ma et al., 2009; Park et al., 2009).

Since then, the PYR/PYL-PP2C interaction has been utilized to construct FRET-based biosensors to study the localisation of ABA in plants (Jones et al., 2014; Waadt et al., 2014). Furthermore, ABA-induced dimerization has been utilized as a genetic switch to regulate gene expression in mammalian and yeast cells (Cunningham-Bryant et al., 2019; Gao et al., 2016; Liang et al., 2011). These genetic switches are based on a Y2H architecture, using two fusion proteins (Figure 1B). A PYR/PYL receptor is fused to an activation domain (AD), and a PP2C is fused to a DNA binding domain (BD). In the presence of the inducer ABA, the fusion proteins dimerize at the promoter and the gene of interest is being expressed. This design solely relies on the ABA-induced dimerization and does not require a functional phosphatase. In a recent study, a highly sensitive ABA biosensor for use in mammalian cells was developed based on the same design principle (Kim et al., 2022). In addition, two latest studies changed the ligand specificity of PYR/PYL receptors and developed PYR/PYL-PP2C-based biosensors for various non-native ligands, including herbicides, cannabinoids and organophosphates (Beltrán et al., 2022; Zimran et al., 2022).

So far, none of these FRET or transcription factor-based biosensors have been used for screening enzyme libraries or other metabolic engineering applications. Still, the studies provide valuable information about *in vivo* ABA sensing and blueprints for biosensor designs. All the mentioned biosensors are based on ABA-induced formation of heterodimers; however, different PYR/PYL variants (e.g. PYR1, PYLcs) and PP2C variants (e.g. ABIcs, HAB1) have been used (Beltrán et al., 2022; Liang et al., 2011). Moreover, various ADs (e.g. Gal4AD, VPR) and BDs (e.g. Gal4BD, dCas9) have been tested (Gao et al., 2016; Zimran et al., 2022). Different reporter genes were used (e.g. luciferase, fluorescent proteins) and the sensors were characterised in different hosts (Liang et al., 2011; Zimran et al., 2022). The studies report different dynamic and operational ranges. To find a suitable candidate for HTS of BcABA1 and BcABA2 mutants, different biosensor designs need to be compared and characterised in yeast cells and the effects of *in vivo* ABA production on the biosensor response curve needs to be tested.

Enzyme engineering is usually an iterative process with multiple rounds of screening, validation and successive combination of the identified beneficial mutants. Having a dedicated platform strain which is specifically optimised for screening applications, and which can be adapted in future screening rounds with minimal genetic adjustments is useful for streamlining this process and increasing its throughput. Various examples of screening platforms have been published already (Kang et al., 2017; Liu et al., 2021; Scott et al., 2022; Shan et al., 2020). Platform strains are especially advantageous for multi-step product pathways like ABA biosynthesis, where several heterologous enzymes likely need to be optimised to increase the metabolic flux to the product. In this study, we combine synthetic biology approaches to develop HTS platform strains with tuneable ABA production (Figure 1C).

## Materials and Methods

### Molecular Cloning

PCR reactions were performed using PrimeSTAR HS DNA Polymerase (Clontech) or Phusion High-Fidelity DNA polymerase (Thermo Fisher Scientific). Primer sequences (ordered from Eurofins Genomics or Integrated DNA Technologies) are listed in Supplementary Table S1 and PCR conditions were set according to the manufacturer’s instructions. Details about individual PCR reactions are listed in Supplementary Table S2. GeneJet Kits (Thermo Fisher Scientific) were used for PCR product purification and minipreps. PCR products and plasmids were verified by Sanger DNA sequencing (performed by Eurofins Genomics). Yeast-codon-optimized *A. thaliana* genes were ordered from GenScript Biotech Corp. The *yeGFP* sequence published by Cormack *et al*. (1997) was used. The yeast Modular Cloning (MoClo) workflow was used to construct the episomal plasmids and plasmids containing genomic integration cassettes (Engler et al., 2008; Lee et al., 2015; Otto et al., 2021), with the exception of pCfB2904-CPR1 and pCfB2909-ABA4 for which Gibson assembly (Gibson et al., 2009) was used. For the Gibson assembly of pCfB2904-CPR1 PCRs 176/197 and 185/198 were used. For the Gibson assembly of pCfB2909-ABA4 PCRs 175/178 and 176/179 were used. Plasmids used for yeast transformations are listed in Table 1. MoClo assemblies are listed in Supplementary Table S3.

**Table 1:**
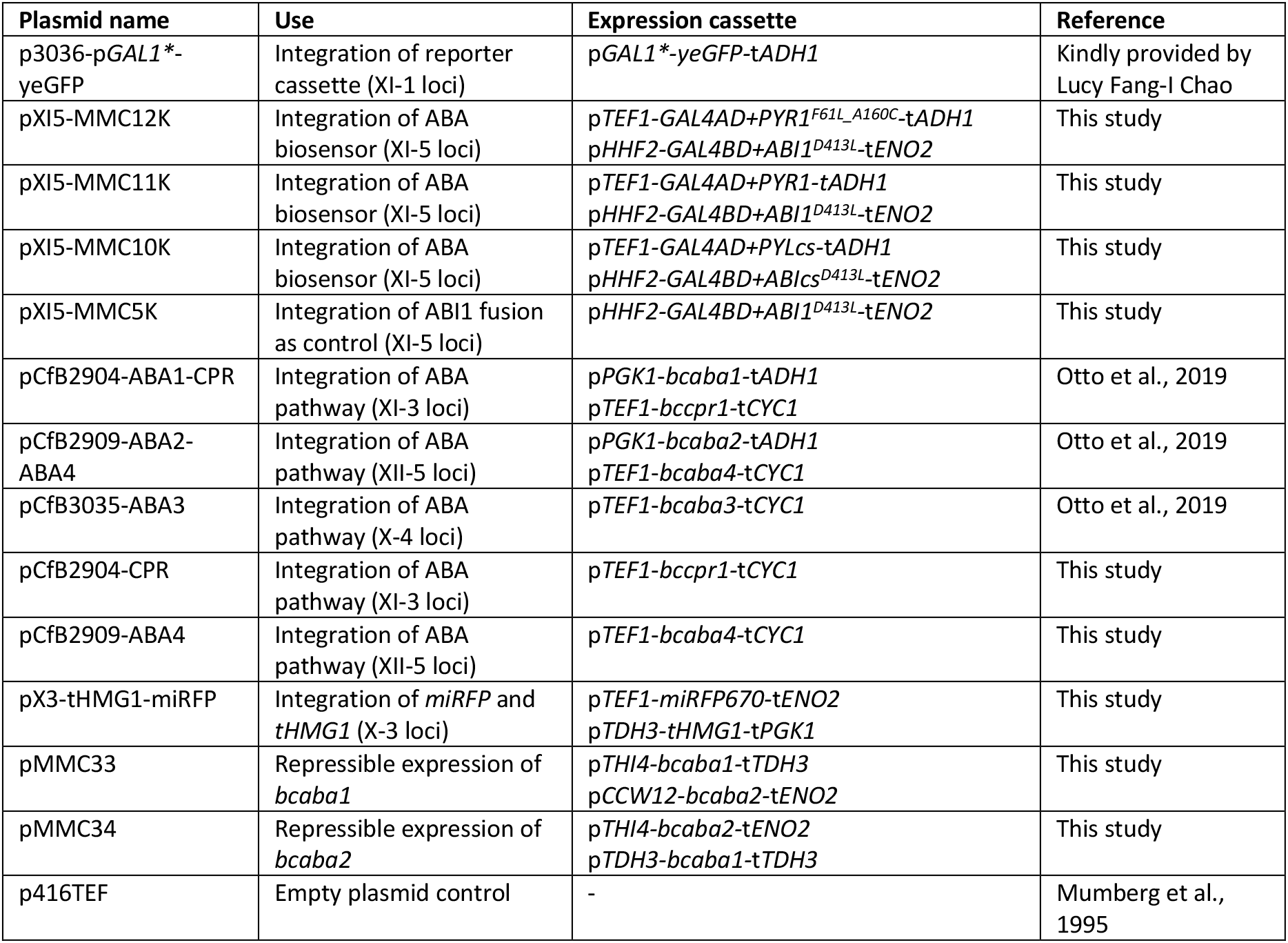
Plasmids used for yeast transformations.

The vectors pGBDU-C1 and pGBAU-C1 (James et al., 1996) were used to amplify *GAL4AD* and *GAL4BD* (Supp. Table S2). ABA biosensor fusion proteins were constructed within the MoClo framework by using type-3a and type-3b parts for the yeast and plant proteins respectively (Supp. Table S3). This results in a Gly-Ser linker between the two proteins. The yeast proteins are located at the N-terminus, the plant proteins are located at the C-terminus of the translated protein.

### Microorganisms and media

NEB® 5-alpha Competent *E. coli* cells (New England Biolabs) were used for plasmid amplification. *E. coli* was grown at 37 °C in liquid Luria-Bertani (LB) media or cultivated stationary using LB agar plates with appropriate antibiotics.

*S. cerevisiae* strains used in this study are listed in Table 2 and displayed in Figure 2. Yeast strains were grown at 30 °C in liquid yeast extract peptone dextrose (YPD) media, buffered synthetic defined (SD) media (Prins and Billerbeck, 2021, pH 6, 0.5x, without thiamine, without uracil) or mineral media supplemented with uracil and histidine (adapted from (Verduyn et al., 1992)) shaking at 220 rpm. For cultivation on agar plates YPD media or SD media (unbuffered, without uracil) was used. For all yeast cultivations 20 g/L glucose was used. Supplementary Table S4 lists detailed media compositions. Mineral media was used for the biosensor characterisation (Figure 3). SD media was used for experiments involving p*THI4*-regulation of *bcaba1* or *bcaba2* (Figure 4, Figure 5).

**Figure 2:**
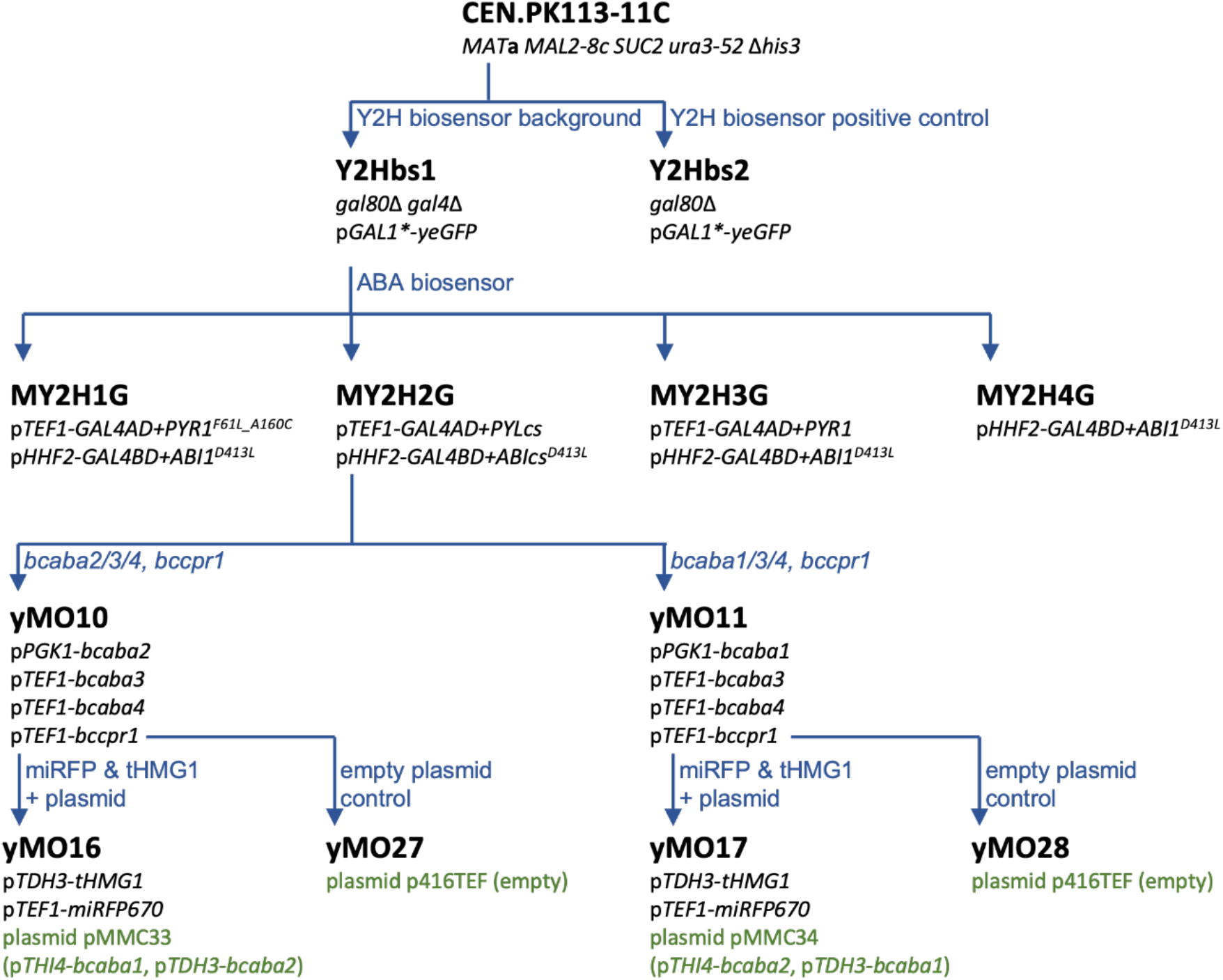
Pedigree tree of strains constructed in this study. Plasmids and their expression cassettes are shown in green. More detailed information can be found in Table 1 and Table 2.

**Figure 3:**
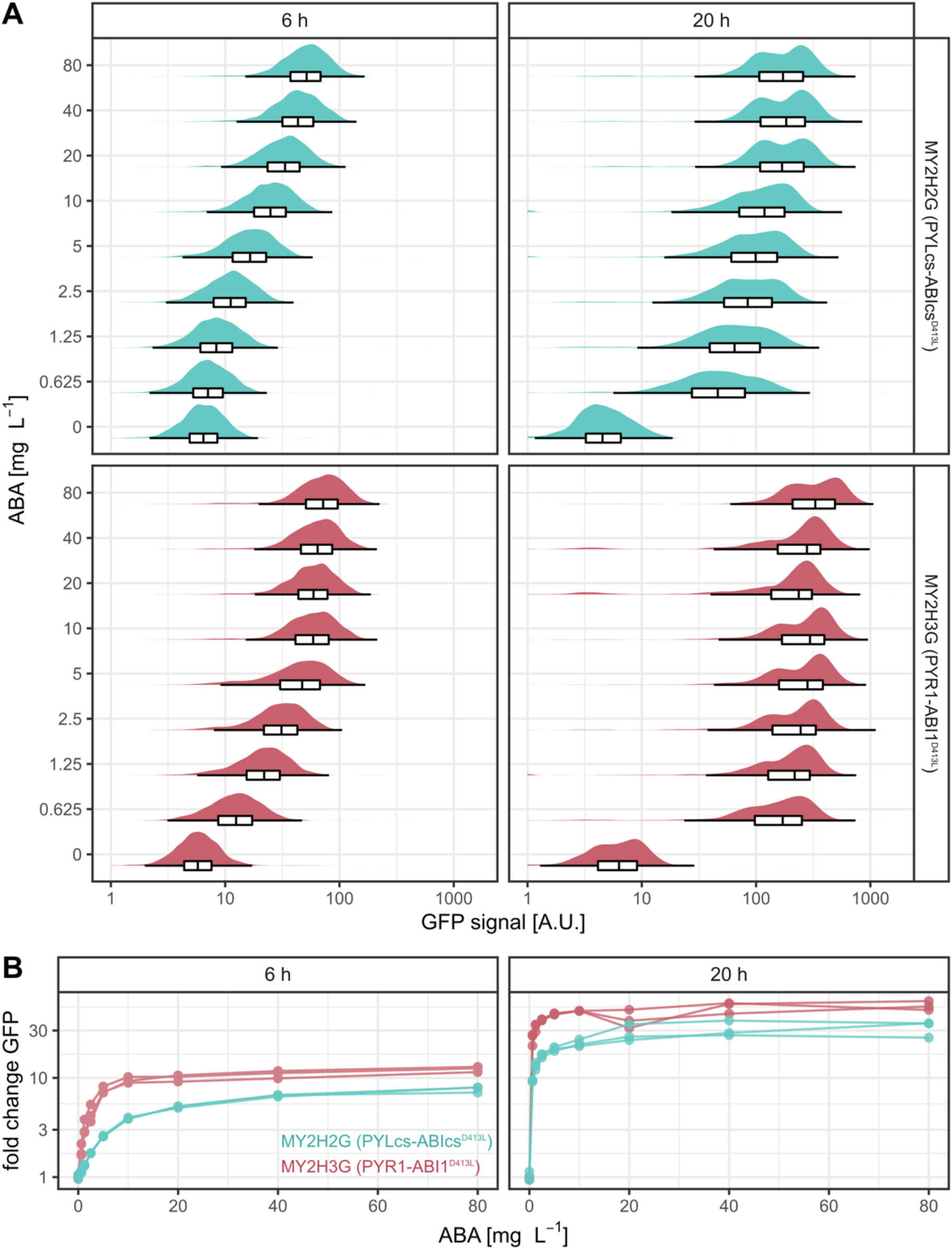
Characterisation of two Y2H ABA biosensor candidates. MY2H2G (PYLcs-ABIcs^D413L^, teal) and MY2H3G (PYR1-ABI1^D413L^, red) were induced with different concentrations of ABA and grown in mineral media. Flow cytometry was performed after 6 h and 20 h of incubation. **A:** Overlayed histograms and boxplots (in the style of Tukey, outliers not shown) showing GFP signal for different ABA concentrations. For readability only one representative replicate is shown. Supplementary Figure S3 and S4 contain additional replicates as well as positive and negative controls. **B:** ABA biosensor response curves obtained from the data shown in A and Supplementary Figure S3 and S4. The fold change was determined by normalising the median GFP signals to the sample without ABA of the same strain. Three biological replicates are shown per strain.

**Figure 4:**
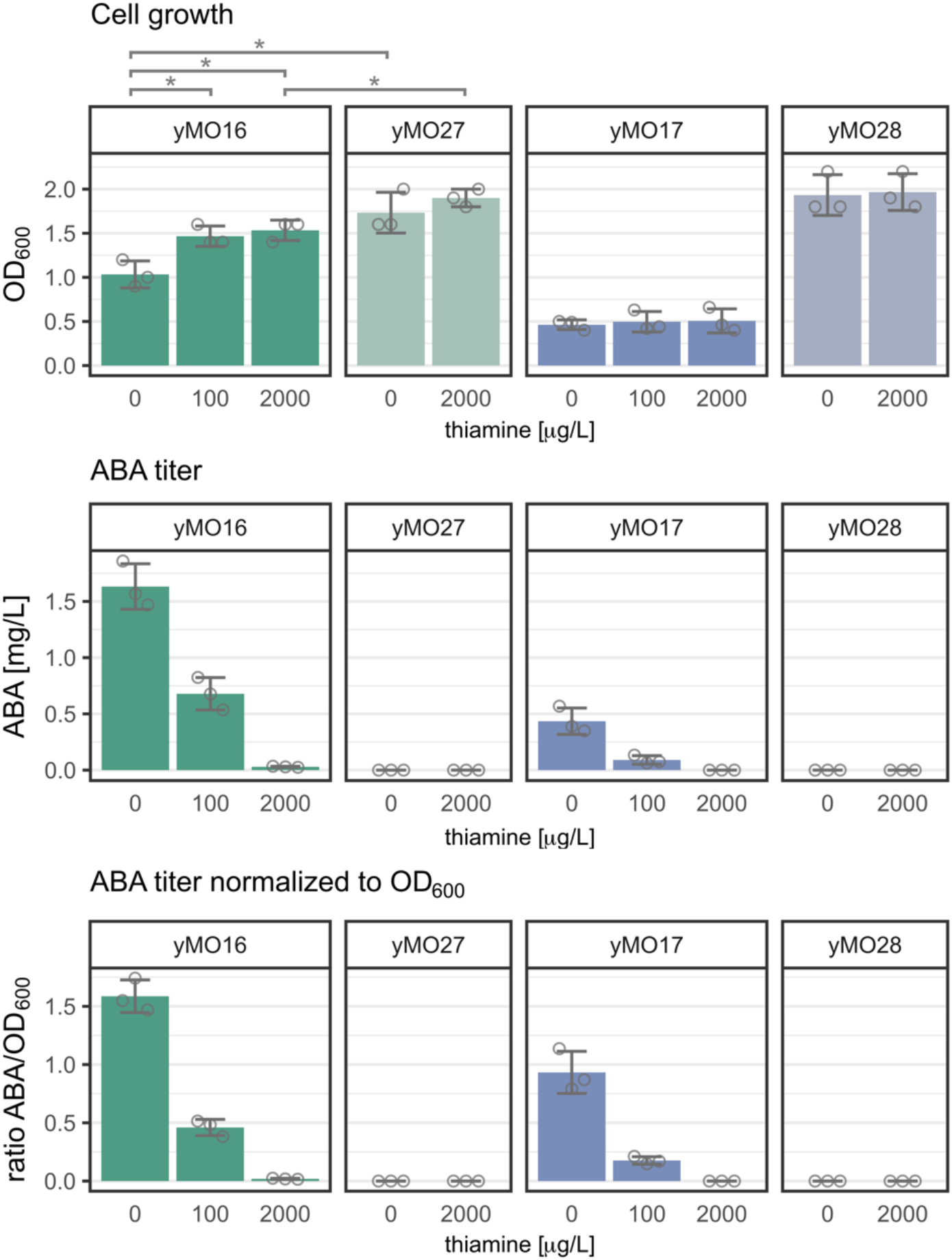
Effect of thiamine concentration on cell growth and ABA production in strains with repressible *bcaba1* or *bcaba2* expression and control strains. Mean and standard deviation were calculated using three biological replicates. Individual measurements of the replicates are included as grey rings. The strains yMO16 (p*THI4-bcaba1*, dark green), yMO17 (p*THI4-bcaba2*, dark blue), yMO27 (light green, lacking *bcaba1*) and yMO28 (light green, lacking *bcaba2*) were cultivated for 20 h in buffered SD media (pH 6) with different thiamine concentrations. Afterwards, the OD_600_ was measured, the ABA concentration was determined in the culture supernatant. No ABA was detected for yMO17 in the 2000 µg/L thiamine condition or for the control strains yMO27 and yMO28 in any condition. To reduce sample size the control strains were cultivated only in the absence of thiamine and in the maximal concentration that was used. When not apparent, significance (Student’s t-test, two-sided) is displayed as “*” for p<0.05.

**Figure 5:**
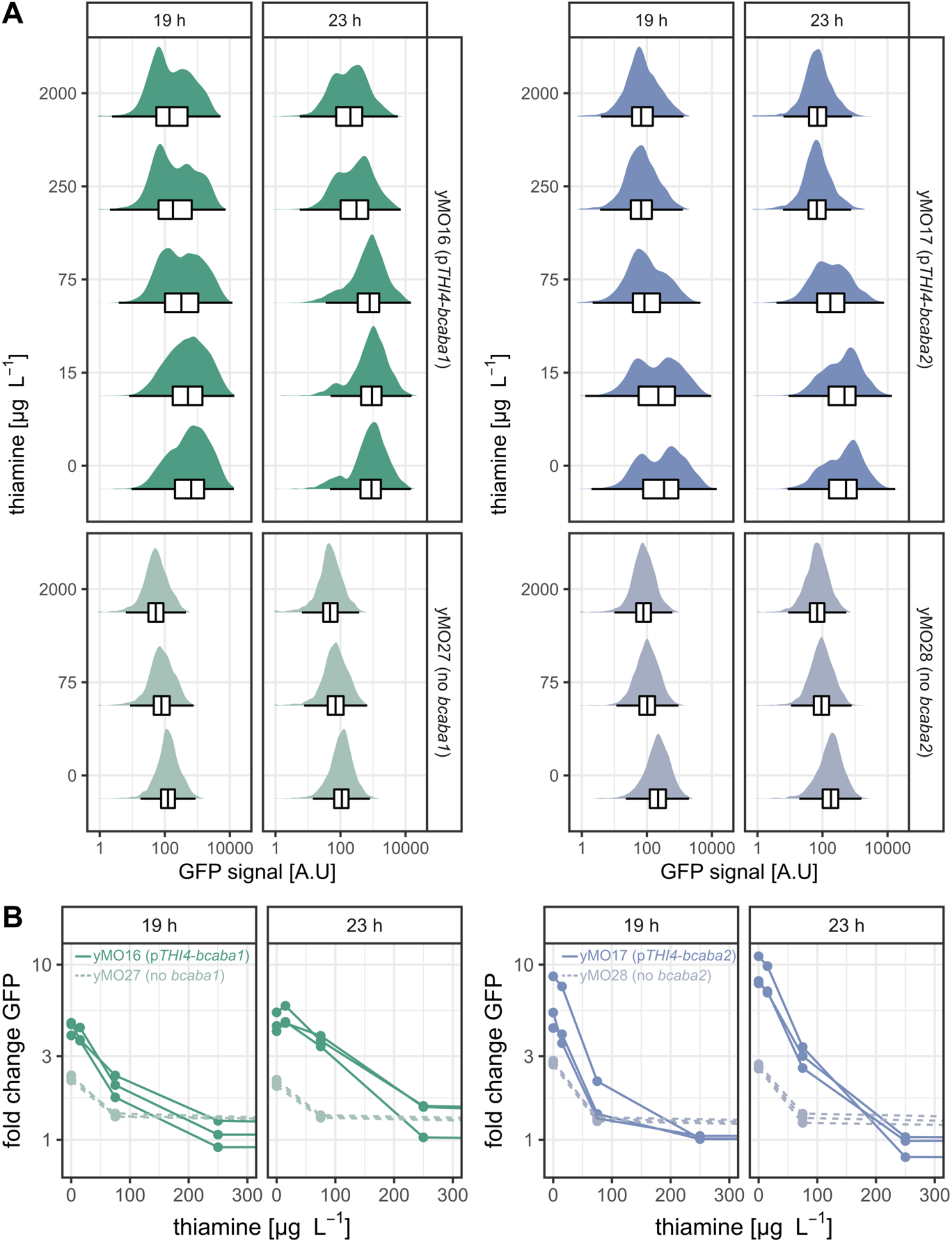
ABA biosensor response in varying thiamine concentrations. The genes *bcaba1* (yMO16, dark green) or *bcaba2* (yMO17, dark blue) are regulated by the thiamine-repressible promoter p*THI4*. The negative controls yMO27 (light green) and yMO28 (light blue) are unable to produce ABA. Strains were cultivated in buffered SD media (pH 6) containing different thiamine concentrations and flow cytometry analysis was performed after 19 and 23 h. **A:** Overlayed histograms and boxplots (in the style of Tukey, outliers not shown) showing GFP signal for different thiamine concentrations. For readability only one representative replicate is shown. Supplementary Figure S9 and S10 show all replicates as well as the positive control expressing wild type *GAL4*. **B:** Fold change GFP reporter output in relation to thiamine concentration in the media obtained from the data shown in A and Supplementary Figure S9 and S10. Dashed lines were used for the control strains yMO27 and yMO28. For readability the x-scale was cropped at 300 µg/L thiamine. The fold-change was determined by normalising the median GFP signals to the 2000 µg/L thiamine sample of the same strain. Three biological replicates are shown per strain.

**Table 2:**
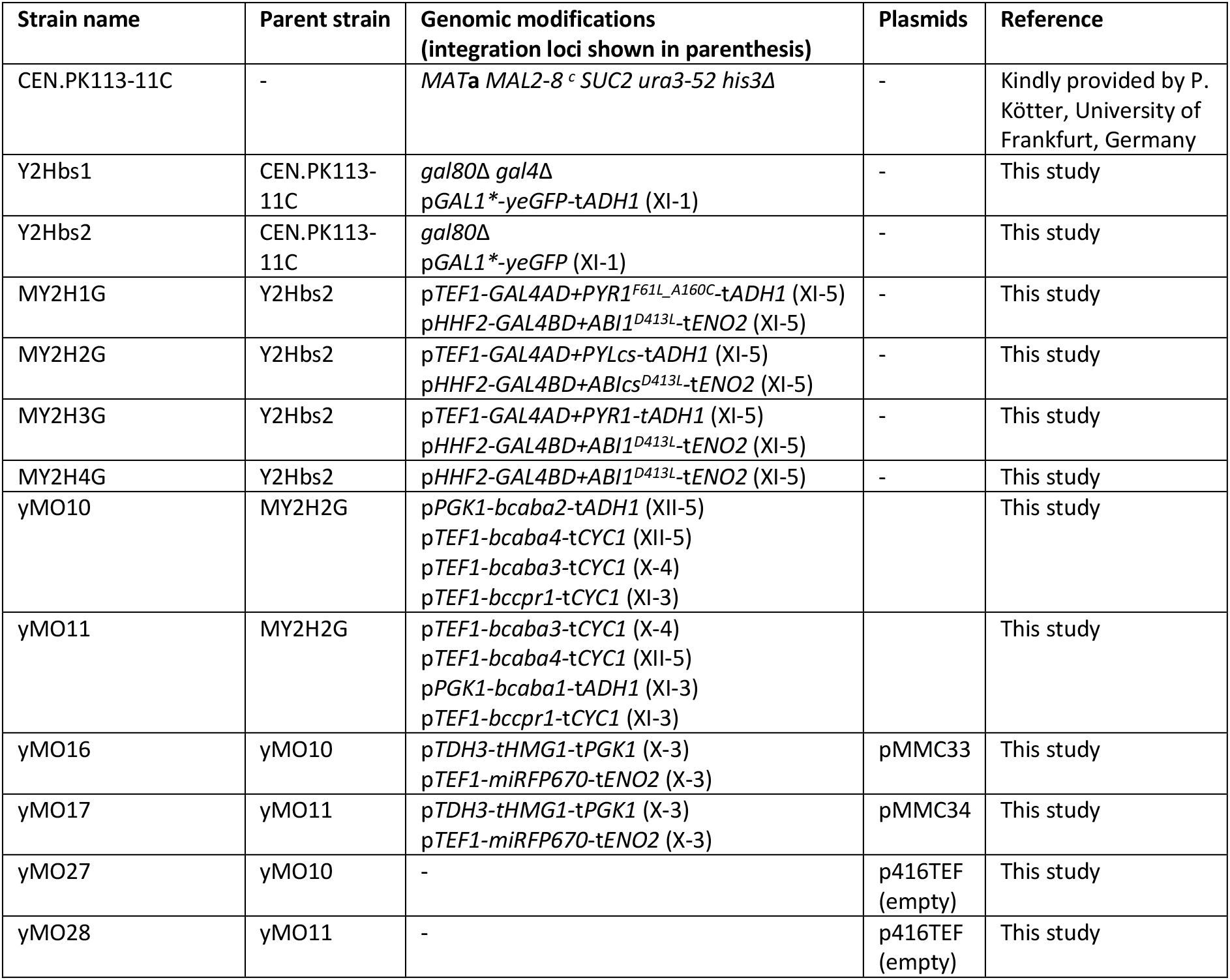
*S. cerevisiae* strains used in this study.

### Strain construction

For genomic integrations the Cas9 and gRNA helper plasmids from the EasyClone-MarkerFree toolkit were used and the protocol was followed as described (Jessop-Fabre et al., 2016). The *GAL4* and *GAL80* deletion were performed following the protocol of Mans et al. (2015). For gRNA expression the plasmid pMEL10 (Mans et al., 2015) was used. Supplementary Table S5 lists the gRNA and repair oligo sequences. Genetic modifications were verified by colony PCR using primers published previously (Jessop-Fabre et al., 2016; Otto et al., 2021) or primers provided by the Yeastriction tool (Mans et al., 2015). SapphireAmp polymerase (Takara Bio) was used for yeast colony PCRs.

### Flow cytometer analysis

Precultures were grown in mineral media (Figure 3) or buffered SD media supplemented with 2000 µg/L thiamine (thiamine HCL, Sigma-Aldrich) (Figure 5) for 48 h. For the experiment displayed in Figure 5, the precultures were centrifuged (5 min, 1500 xg) and the cell pellet was washed with milliQ-H_2_O to remove the remaining thiamine. Main cultures were inoculated at OD_600_ 0.1 (96-half-deepwell microplates, 250 µL culture volume, transparent bottom, EnzyScreen). Cells were grown in varying concentrations of (S)-(+)-ABA standard (>98%, Cayman Chemicals) (Figure 3) or varying concentrations of thiamine (Figure 5). At the indicated time points cell cultures were diluted in milliQ-H_2_O and flow cytometry analysis was performed using a Guava® easyCyte^TM^ HT System. FSC and SSC parameters were used to gate yeast cells and remove very large and very small particles. A FSC-A/FSC-H plot was used to discriminate doublets. For strains with simultaneous *yeGFP* and *miRFP* expression the fluorescence compensation wizard was used in the FlowJo software (BD Life Sciences, version 10.8.1).

### ABA extraction

Precultures were grown in buffered SD media supplemented with 2000 µg/L thiamine (thiamine HCL, Sigma-Aldrich) for 48 h. The precultures were centrifuged (5 min, 1500 xg) and the cell pellet was washed with milliQ-H_2_O to remove the remaining thiamine. Main cultures were inoculated at OD_600_ 0.1 in buffered SD media with varying thiamine concentrations (24-deepwell microplates, 2.5 mL culture volume, square wells, pyramid-bottom, EnzyScreen). After 20 h, the OD_600_ was measured, the cultures were centrifuged (5 min, 1500 xg) and 1 mL supernatant was transferred to 2-mL Eppendorf tubes. 1 mL of ethyl acetate (>99.9% Sigma-Aldrich) and 0.5% (v/v) formic acid (>98%, Sigma-Aldrich) was added to the 2-mL Eppendorf tubes containing 1 mL culture supernatant. The tubes were vortexed for 10 s, then centrifuged (10 min, 13.000 xg, 4°C) and 0.8 mL of the supernatant was transferred to a new Eppendorf tube. The solvent was evaporated using a Genevac miVac (45 min, 20 mBar, 45 °C). The pellet was reconstituted in 0.8 mL methanol (>99.9%, Sigma-Aldrich), centrifuged (10 min, 13.000 xg, 4 °C) and ≈0.5 mL of the supernatant was transferred to an HPLC (high-performance liquid chromatography) vial.

### ABA quantification

The samples were analysed in an Agilent 6120 Single Quadrupole mass spectrometer (MS) with an Agilent Infinity 1260 HPLC system consisting of a binary pump, autosampler and thermostat column compartment. Ionization was performed using an atmospheric pressure electrospray ionization (API-ES) source (positive mode). Compounds were separated on an Agilent Poroshell 120 EC-C18 (2.7 μm, 3.0 × 50 mm) column (maintained at 40 °C) using a water-acetonitrile gradient with 0.04% formic acid in both solvents. The gradient started with 95% water and, over 5 min, gradually changed to 95% acetonitrile (>99.5%, Sigma-Aldrich). After a 2 min hold, the gradient was ramped back to 95% water over 3 min. The sample injection volume was set to 10 μL. ABA was quantified by selected ion monitoring (m/z = 265) and (S)-(+)-ABA standard (>98%, Cayman Chemicals) was used to fit a calibration curve (2^nd^ polynomial).

### Growth Profiler analysis

A Growth Profiler 960 (EnzyScreen) was used for continuous growth monitoring in 96-half-deepwell microplates (transparent bottom, EnzyScreen). The culture volume was 250 μL per well and pictures were taken every 30 min.

### Software

DNA constructs were design and analysed using Benchling (www.benchling.com). FlowJo (BD Life Sciences, version 10.8.1) and R Studio (RStudio Team, 2022) were used to analyse and plot the data. Relevant R packages include ggplot2 (Wickham, 2016), tidyverse (Wickham et al., 2019) and growthrates (Petzoldt, 2022).

## Results and Discussion

### Characterisation of ABA biosensor candidates in feeding experiments

PYR/PYL ABA receptors and plant PP2Cs have been utilized in basic research to elucidate ABA sensing and transport in plants, and have further been applied in synthetic biology to regulate gene expression (Cunningham-Bryant et al., 2019; Jones et al., 2014; Liang et al., 2011; Ma et al., 2009; Park et al., 2009; Waadt et al., 2014). None of the PYR/PYL-based constructs in the literature have been utilized for metabolic engineering purposes until now. The desired biosensor characteristics, such as the sensor’s response curve and the reporter gene used, are highly application dependent. A large dynamic range is desirable for most purposes; however, HT-screening applications also require a large operational range. The operational range needs to be compatible with the current production capabilities, with the current titer being at its lower end and the desired titer at its higher end. An ABA biosensor designed for yeast cell factories would allow HTS for improved ABA pathway enzymes.

We decided to use a Y2H-based design (Liang et al., 2011) for the biosensor instead of a dCas9-based design (Cunningham-Bryant et al., 2019; Gao et al., 2016) for two reasons: firstly the published dCas9-based activator appears to be too sensitive for our current production strain, and secondly the presence of dCas9 and its cognate gRNA would hamper subsequent Cas9-mediated genome engineering, potentially requiring the removal of the biosensor before the engineering step and re-introduction afterwards. For a direct comparison, three different PYR/PYL-PP2C combinations with varying ABA affinities were chosen from the literature. Each candidate consists of two hybrid proteins: the Gal4AD fused to a variant of an *A. thaliana* PYR/PYL ABA receptor and the Gal4BD fused to a variant of the *A. thaliana* ABI1 phosphatase (Figure 1B). In all used ABI1 variants, the phosphatase activity was deactivated to increase orthogonality of the biosensor (the cross-reactivity between the native yeast metabolism and the biosensor setup). The three sensor candidates are listed and described in Table 3.

**Table 3:** ABA biosensor candidates analysed in this study.

| Gal4AD fusion | Gal4BD fusion | Expected ABA affinity | Notes |
| --- | --- | --- | --- |
| PYR1<sup>F61L_A160C</sup><br>(high sensitivity) | ABI1<sup>D413L</sup><br>(catalytically inactive) | High | PYR1<sup>F61L_A160C</sup> was previously identified in a Y2H β-galactosidase assay (Elzinga et al., 2019), the D413L mutation in ABI1 was previously used in a FRET-based sensor (Waadt et al., 2014) |
| PYLcs<br>(truncated) | ABIcs<sup>D413L</sup><br>(truncated) | Low | PYL1 and ABI1 were truncated and reduced to the interacting complementary surfaces (cs) (Liang et al., 2011), the design was used in mammalian cells and yeast to regulate gene expression using ABA as an inducer (Cunningham-Bryant et al., 2019; Gao et al., 2016) |
| Wild type PYR1 | ABI1<sup>D413L</sup><br>(catalytically inactive) | Medium | The D413L mutation in ABI1 was previously used in a FRET-based sensor (Waadt et al., 2014) |

A p*GAL1*-yeGFP* reporter cassette was integrated in the genome of the sensor background strain. The native *GAL1* promoter contains two binding sites for the transcription factor Mig1, which represses various catabolic genes in the presence of glucose (Frolova et al., 1999; Nehlin et al., 1991; Schüller, 2003). To avoid glucose-dependent effects on reporter gene expression and increase the biosensor’s orthogonality further, the Mig1 binding sites were removed from p*GAL1*, resulting in p*GAL1\**. Furthermore, the native *GAL4* and *GAL80* genes were deleted in the background strain to prevent reporter gene activation by the wild type Gal4 activator and Gal80-mediated repression of the reporter in the absence of galactose.

In this background strain, two strong promoters with similar expression levels (Lee et al., 2015) were used to express the two fusion proteins, p*TEF1* was used for the Gal4AD fusion and p*HHF2* was used for the Gal4BD fusion. After inoculation of the main cultures, the strains were induced with different concentrations of ABA and later analysed using a flow cytometer.

Initially, the sensor proteins were expressed from a centromeric plasmid. However, the analysis showed large cell-to-cell variability in all three candidate strains (Supp. Fig. S1), likely due to cells losing the plasmid and/or cells carrying more than one copy of the plasmid (Singh and Anthony Weil, 2002). From this analysis, it was also apparent that the PYR^F61L_A160C^ variant was likely too sensitive to be useful for the ABA cell factory. For the PYR^F61L_A160C^ variant, 5 mg/L ABA resulted in virtually maximal induction of the biosensor (Supp. Fig. S1). In addition, when no ABA was added to the culture, the PYR^F61L_A160C^ variant showed the leakiest GFP expression of the three tested candidates. We therefore decided to omit the PYR^F61L_A160C^-based sensor candidate in following experiments.

For the subsequent analysis, the sensor candidates were integrated into the genome to facilitate stable gene expression. Strain MY2H2G contains the PYLcs-ABIcs^D413L^ combination and strain MY2H3G contains the PYR1-ABI1^D413L^ combination (Figure 2). After addition of ABA in various concentrations (0.625 mg/L to 80 mg/L; 2.4 µM to 302.7 µM), the strains were analysed during the glucose phase, at 6 h of cultivation, and after the diauxic shift in the ethanol phase, at 20 h of cultivation (see Supp. Fig. S2 for growth profiles). Addition of ABA did not affect cell growth (Supp. Fig. S2). As expected, feeding ABA did not affect the GFP output of the positive control strain Y2Hbs2 which expresses the wild type *GAL4* gene leading to constitutive p*GAL1\** activation and a high GFP signal (Supp. Fig. S3, Supp. Fig S4). Likewise, the low GFP signal of the negative control strain MY2H4G, expressing Gal4BD-ABI1^D413L^ but lacking a Gal4AD fusion protein, did not change in response to ABA (Supp. Fig. S3, Supp. Fig S4). Addition of ABA led to an increase in GFP signal in MY2H2G and MY2H3G visible at 6 h and at 20 h of cultivation (Figure 3). For both strains, at 6 h a gradual increase in GFP signal was visible with increasing ABA concentration, whereas at 20 h the GFP increase was larger but less gradual. Based on the median for the 6 h measurements, the PYR1-ABI1^D413L^ combination reached a near maximal GFP output for 10 mg/L ABA, whereas the PYLcs-ABIcs^D413L^ combination’s signal plateaued around 40 mg/L. For the 20 h measurements, both biosensors reached the maximal signal earlier, at 5 mg/L and 20 mg /L for PYR1-ABI1^D413L^ and PYLcs-ABIcs^D413L^, respectively.

A clear correlation of genotype to phenotype (in this case sensor output) is required for HT screenings and it is therefore paramount to minimise noise that results in cell-to-cell variation. Compared to the strains expressing the sensor proteins from a centromeric plasmid (Supp. Fig. 1) less cell-to-cell variability was visible in the populations of the strains carrying the genomic integrations (Figure 3A). For both strains the populations of the 6 h measurements were more uniform compared to the later measurements. *S. cerevisiae* cells can form distinct populations of budded and un-budded cells that vary in forward scatter (FSC) and side scatter (SSC) (Ottoz et al., 2014). We observed a correlation between GFP and SSC/FSC (Supp. Figure S5), indicating that budded cells, which scatter more light, exhibit a higher GFP signal. These cells likely had more time to accumulate GFP than the younger and smaller un-budded cells. The difference appeared to be more pronounced after 20 h, resulting in more population heterogeneity. However, in library screening experiments this effect can be taken into account by gating budded and un-budded populations separately.

The relative ABA affinities of the biosensor candidates coincided with the literature (Elzinga et al., 2019; Jones et al., 2014; Waadt et al., 2014), with the PYR1-ABI1^D413L^ biosensor being more sensitive to ABA compared to the PYLcs-ABIcs^D413L^ biosensor. Only a marginal change in GFP output was observable for the PYLcs-ABIcs^D413L^ combination with 0 to 1.25 mg/L ABA at 6 h. For the PYR1-ABI1^D413L^ biosensor, a clear increase in GFP signal was visible for the same conditions.

The tested biosensor candidates differed in dynamic and operational range and both features were dependent on the cultivation time (Figure 3B, summarised in Table 4). For 6 h of cultivation, the dynamic range was ≈8-fold for PYLcs-ABIcs^D413L^ and ≈12-fold for PYR1-ABI1^D413L^. For 20 h of cultivation, the dynamic range was ≈32-fold and ≈54-fold, respectively. The effective operational range for PYLcs-ABIcs^D413L^ at 6 h extended from 2.5 to 80 mg/L ABA (9.5 to 302.7 µM). The effective operational range for PYR1-ABI1^D413L^ at 6 h extended from 0.625 to 10 mg/L ABA (2.4 to 37.8 µM). The operational ranges at 20 h were 0.625 mg/L to 20 mg/L (2.4 – 75.7 µM), and 0.625 mg/L to 5 mg/L (2.4 – 18.9 µM) for PYLcs-ABIcs^D413L^ and PYR1-ABI1^D413L^, respectively. However, 0.625 mg/L ABA was the lowest concentration tested and lower concentrations are likely detectable at this timepoint. Different sensor designs, reporter setups, incubation times and host organisms make direct comparisons with other studies difficult. Nonetheless, a similar µM operational range has been reported for a PYLcs-ABIcs design in mammalian cells using a luciferase assay as output (Liang et al., 2011).

**Table 4:**
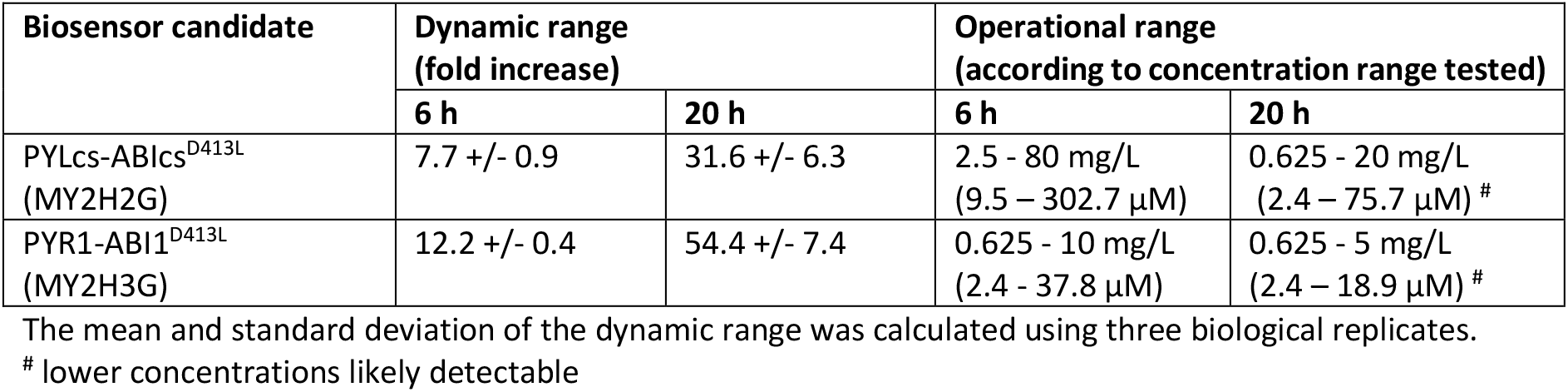
Dynamic and operational range of biosensor candidates.

The current proof-of-concept ABA cell factory produced 5 to 10 mg/L ABA (Otto et al., 2019). While the dynamic range of PYR1-ABI1^D413L^ was higher compared to PYLcs-ABIcs^D413L^, the operational range of PYLcs-ABIcs^D413L^ was more suitable for the product titer of the ABA-producing strain, with sufficient range to detect and distinguish ABA overproducers. We therefore chose to use the PYLcs-ABIcs^D413L^ sensor strain MY2H2G for further analysis.

In this ABA feeding experiment, analysis after 6 h of cultivation was preferable to the 20 h timepoint with regards to the operational range. In contrast to the immediate induction at the beginning of this experiment, ABA accumulated over several hours (peaking after ≈ 16 h) in the culture of ABA-producing strains (Otto et al., 2019). In ABA-producing strains the biosensor response is likely shifted.

### Design of BcABA1 or BcABA2 screening strains

Preceding this study, we demonstrated that the two CYP enzymes, BcABA1 and BcABA2 catalyse limiting steps in the ABA biosynthetic pathway in yeast (Otto et al., 2019). We therefore aim to screen for improved variants of the two enzymes to increase the productivity of ABA cell factories.

The success of HTSs is highly dependent on having a biosensor with optimal characteristics. Once an improved enzyme is found the biosensor usually needs to be fine-tuned, e.g. its operational range needs to be adjusted to the increased product titer. However, fine-tuning biosensors remains challenging and time-consuming. Instead of further optimising the PYLcs-ABIcs^D413L^ biosensor we decided to construct a screening strain in which ABA production can be regulated by fine-tuning the expression levels of the enzyme that is to be engineered. This approach provides several potential advantages: (1) the full operational range of the biosensor can be utilized by adjusting the ABA titer; (2) the screening selectivity can be increased by adjusting ABA production “on the fly” between FACS enrichment rounds, after the initial analysis and/or first sorting of the library; (3) ABA production can be turned off or minimized allowing for a negative selection step, in which false positives (with ABA-independent high GFP signal) can be removed; and (4) the same biosensor design can be used in subsequent screenings when an improved enzyme should be engineered further. To achieve this, the native, thiamine-repressible promoter p*THI4* was used to regulate *bcaba1* or *bcaba2* expression in the screening strain (Figure 1C). The *THI4* promoter was shown to have a high dynamic range of ≈1000-fold induction and low leakiness in repressive conditions (Rantasalo et al., 2018).

MY2H2G was used as a background strain and two screening platform strains were constructed, one for each of the CYPs in the ABA pathway, BcABA1 and BcABA2 (Figure 2). Both strains constitutively express *bcaba3*, *bcaba4* and the *B. cinerea* CPR gene *bccpr1*. The BcABA1 screening strain, yMO16, carries the repressible p*THI4*-*bcaba1* expression cassette, while the BcABA2 screening strain, yMO17 carries the repressible p*THI4*-*bcaba2* expression cassette.

In screenings, it is crucial that the enzyme that is being engineered is the rate-limiting step in the production pathway. To eliminate current and potential future bottlenecks in the screening strains, two copies of *bcaba2* are being expressed in yMO16 and two copies of *bcaba1* are being expressed in yMO17. While the supply of the precursor FPP is not limiting ABA production when using the wild type ABA pathway genes (Otto et al., 2019), we nonetheless decided to alleviate a potential future bottleneck by expressing a truncated version of the *HMG1* gene (*tHMG1*) (Donald et al., 1997; Polakowski et al., 1998). Hmg1 catalyses the conversion of 3-hydroxy-3-methyl-glutaryl-CoA to mevalonate in the mevalonate pathway, the universal precursor pathway for terpenoids in yeast. Expression of *tHMG1* is a commonly used strategy to increase metabolic flux through the mevalonate pathway (Ro et al., 2006; Scalcinati et al., 2012).

Extrinsic noise, originating from e.g. cell size, availability of RNA polymerases and cell cycle stage, affect overall expression levels and is a major contributor to population heterogeneity (Raser and O’Shea, 2004; Swain et al., 2002). Extrinsic noise will affect the biosensor reporter output and make screening more difficult. Furthermore, hyperactive enzyme variants might cause growth defects that could influence overall protein expression levels, affecting expression of the mutant gene and the reporter output. Constitutive expression of a second fluorescent proteins as a “noise reporter” has been used to normalize a reporter output to extrinsic noise, making it possible to distinguish increased GFP signal caused by higher inducer concentration from other unrelated factors (Liang et al., 2012; Scott et al., 2022). For normalization purposes, yMO16 and yMO17 constitutively express *miRFP670* (Shcherbakova et al., 2016), which is detectable in our flow cytometer and FACS and has low spectral overlap with yeGFP.

Two additional strains were constructed as negative controls. The strains yMO27 and yMO28 (Figure 2) are lacking p*THI4*-*bcaba1* and p*THI4*-*bcaba2* respectively. BcABA1 and BcABA2 are essential for ABA production (Siewers et al., 2006, 2004). Assuming that the biosensor is not responsive to ABA pathway intermediates, the reporter gene should not be activated in the control strains.

### Fine-tuning ABA production via the thiamine-repressible promoter p*THI4*

To test if ABA production can be efficiently regulated by repressing *bcaba1* or *bcaba2*, the strains yMO16 (p*THI4-bcaba1*), yMO17 (p*THI4-bcaba2*), yMO27 (lacking *bcaba1*) and yMO28 (lacking *bcaba2*) were cultivated in varying thiamine concentrations. In a prior report, the thiamine-repressible promoter was tested in a concentration range up to 800 µg/L thiamine, where close to maximal repression was reached (Rantasalo et al., 2018). In this experiment, the analysed strains were cultivated in up to 2000 µg/L thiamine to ensure maximal repression. Strains were precultured in SD media with 2000 µg/L thiamine to minimise the presence of BcABA1 and BcABA2 in the cells before the experiment. After 20 h of main culture, the OD_600_ and ABA titer in the supernatant were measured (Figure 4).

Significant differences in OD_600_ were observed when comparing the different strains. The two negative control strains yMO27 and yMO28 showed the highest cell density and thiamine concentrations did not influence growth in the strains. The p*THI4-bcaba1* containing strain yMO16 had a significantly lower OD_600_ when compared to the control strain lacking *bcaba1*, yMO27, in the same thiamine concentrations. In the absence of thiamine, yMO16 only reached ≈60% of the OD_600_ of yMO27. For 2000 µg/L thiamine, yMO16 reached ≈80% of the yMO27 OD_600_. The p*THI4-bcaba2* strain yMO17 grew substantially worse than any of the other strains, with its OD_600_ being only ≈25% of the strain lacking *bcaba2* yMO28. The OD_600_ measurements showed that expression of the biosensor and the full ABA pathway with additional copies of either *bcaba2* (in yMO16) or *bcaba1* (in yMO17) negatively affected cell growth.

As expected, no ABA was detected in the strains yMO27 and yMO28 lacking *bcaba1* or *bcaba2* respectively. For yMO16 and yMO17 ABA titers decreased with increasing thiamine concentration. Compared to yMO16, significantly less ABA was detected for yMO17 under all conditions. In the absence of thiamine and therefore maximal de-repression of p*THI4*, the ABA titer of yMO17 was about 25% of the yMO16 ABA titer. Miniscule amounts of ABA were detected for yMO16 during growth in 2000 µg/L thiamine, no ABA was detected for yMO17 under the same conditions. When normalized to OD_600_ ABA titers of yMO17 remained significantly lower than yMO16.

Overexpression of heterologous CYPs and CPRs can result in growth defects, likely due to increased production of reactive oxygen species (Paddon et al., 2013; Zangar, 2004). Our data indicate that specifically high expression levels of *bcaba1* are causing a growth defect, as seen in yMO17 with constitutive expression of two copies of *bcaba1* using strong promoters (Figure 4). This is further supported by the fact that yMO16 grew worse in the absence of thiamine. When p*THI4-bcaba1* was de-repressed ABA titer and adaptation to the absence of thiamine did not seem to generally affect cell growth according to the OD_600_ measured after 20 h. However, for heavily engineered strains like yMO16 and yMO17 multiple factors might play a role in the reduced growth besides *bcaba1* expression. Since yMO17 expresses two copies of *bcaba1*, we presume that the reduced ABA titer in this strain compared to yMO16 is due to the growth defect, rather than a bottleneck in the ABA pathway.

The two strains lacking one of the essential enzymes *bcaba1* or *bcaba2*, were expected to accumulate ABA pathway intermediates, specifically ⍺-IE for yMO27 and ⍺-IAA for yMO28 (Figure 1A) (Takino et al., 2019). Chemical standards would be required to identify and accurately quantify the intermediates. Nonetheless, for a general estimate, we analysed the extracted supernatant for compounds with the expected m/z ratio of ⍺-IE (m/z 205 [M+H]) and ⍺-IAA (m/z 235 [M+H]). A peak with the m/z of 205 was not detected in any of the four analysed strains (data not shown). Only the supernatant was analysed and ⍺-IE might either accumulate in the cell, or be degraded or modified by native yeast enzymes. In line with our expectations, a compound with the m/z 235 (expected for ⍺-IAA) at 6.8 min retention time accumulated in yMO28 independent of thiamine concentration (see Supp. Fig. S7 for data and extended discussion). Taken together the data demonstrate that ABA production can be regulated in a dose-dependent manner by cultivating the expressing p*THI4*-*bcaba1* or p*THI4*-*bcaba2* expressing strains in varying thiamine concentrations.

### Biosensor response in strains with tuneable ABA production

The strains yMO16 and yMO17 can both produce and sense ABA. By using the thiamine-repressible promoter p*THI4* for *bcaba1* or *bcaba2* expression, ABA production can be regulated depending on the thiamine concentration present in the media (Figure 4), and the GFP reporter output is expected to vary accordingly (Figure 1C). In the following experiment we aimed to investigate the biosensor response to the varying ABA titers produced in yMO16 and yMO17. In addition, we intend to validate if the biosensor’s full operational range can be covered by regulating the activity of p*THI4*. The ABA-producing strains yMO16, yMO17 and the respective negative controls yMO27 and yMO28 were cultivated in varying thiamine concentrations and flow cytometry analysis was performed after 19 h and 23 h of cultivation. We previously showed that ABA titers differed significantly at 20 h for both producing strains (Figure 4) and initial data indicated that samples with varying thiamine concentrations can be distinguished best after approximately 20 h of cultivation (Supp. Fig. S6). As before, strains were precultured in media with 2000 µg/L thiamine to minimize activation of the ABA biosensor and accumulation of GFP before the experiment.

As was expected, the median GFP signal of yMO16 and yMO17 increased gradually with decreasing thiamine concentration for both timepoints, coinciding with changes in cell populations as visible in the histograms (Figure 5A). The population heterogeneity for the two strains varied in different conditions and different time points. A gradual transition of the histograms is visible when comparing different thiamine concentrations for a single strain at one timepoint. When comparing the same conditions, the median GFP signal of yMO16 was consistently higher in the later, 23 h measurement. A similar behaviour is visible for yMO17, with the exception of the 2000 and 250 µg/L thiamine condition where the GFP signal barely changed between 19 and 23 h. Overall, the GFP signal was higher in yMO16 than in yMO17, as was expected since less ABA was produced in the p*THI4*-*bcaba2* strain (Figure 4).

The GFP signals of the negative control strains yMO28 and yMO29 were nearly identical. Both strains have a distinct peak and their GFP signal did not change in between the two measurement timepoints. Surprisingly, yMO27 and yMO28 also showed an increase in GFP output depending on thiamine concentration, even though they do not produce ABA (Figure 4) and do not carry a heterologous expression cassette with p*THI4* (only the native *THI4* gene promoter in the genome). However, the increase in GFP signal was substantially lower in the control strains compared to the strains with repressible *bcaba1* or *bcaba2* genes and did not coincide with a shift in cell population. We presume that the shift in GFP for the control strains originates from a change in the native metabolism that is not related to ABA intermediates or the *B. cinerea* genes. This effect is most likely a global shift in metabolic activity and not related to p*GAL1\** regulation since a similar trend was visible for the RFP signal in yMO16 and yMO17 as well as for the GFP signal of a positive control strain with wild type *GAL4* expression (Supp. Fig. S8). Thiamine (vitamin B1) is an essential co-factor for various enzymes of the central carbon metabolism (Perli et al., 2020). In the absence of thiamine, Thi4 and Thi5, two enzymes involved in thiamine biosynthesis, are among the most highly expressed proteins in yeast (Muller et al., 1999). The high cellular demand for Thi4 and Thi5 derives from the fact that both are suicide enzymes that can catalyse only a single turnover (Chatterjee et al., 2011; Coquille et al., 2012; Lai et al., 2012). To the best of our knowledge the effects of lack of thiamine on overall protein expression has not been studied thoroughly enough to explain our observations. It is however conceivable that upregulation in transcription, translation and folding machinery in the cell to meet the demands for thiamine biosynthesis enzymes also affects GFP and RFP expression levels.

The GFP signal of yMO16 was higher compared to the control strain yMO27 for all conditions and at both timepoints, confirming that ABA is being produced in sufficient amounts to trigger the biosensor even at maximal repression in the 2000 µg/L thiamine condition (Figure 5A). For yMO17 the GFP signal for the 250 and 2000 µg/L thiamine conditions were very similar to the negative control yMO28, with slightly more population heterogeneity for yMO17, indicating that the biosensor was not triggered to a meaningful degree under these conditions. These data confirm the previous analysis (Figure 4) where no detectable amounts of ABA were present for yMO17 in 2000 µg/L thiamine.

The flow cytometry analysis was used to quantify the relationship between sensor response and thiamine concentration (Figure 5B). For yMO16, a ≈4-fold induction of the median GFP signal was visible for both the 19 h and 23 h measurements. For yMO17 a ≈6-fold induction and ≈9-fold induction was observed at 19 h and 23 h respectively, which was slightly higher than the dynamic range of the biosensor measured in ABA feeding experiments (Figure 3B, Table 4). The fold increases of the negative controls were smaller with ≈2.2-fold and ≈2.7-fold for yMO27 and yMO28, respectively, and virtually no differences between the measurement timepoints.

As seen in the previous experiment (Figure 3), the ABA biosensoŕs dynamic and operational range varies depending on the time of measurement. The measurements from yMO17 (Figure 5B) indicate that, by modulating the thiamine concentration, it is possible to cover a large section, if not all of the operational range of the ABA biosensor at that point in time. Nonetheless, exact comparisons are difficult, in part because of the thiamine-related effect on the GFP signal visible in the control strains. The data (Figure 5A) confirm that ABA titers in yMO16 were substantially higher after 19 and 23 h of cultivation than in yMO17. We presume that measuring the yMO16 GFP signal at an earlier timepoint when the ABA titer is lower would result in a sensor output similar to what was observed for yMO17, where the samples with high thiamine concentrations were close to the baseline GFP of the control strain. In turn, this would also result in a higher fold-induction, as seen in yMO17 (Figure 5B).

Plants produce ABA via a different biosynthetic pathway than *B. cinerea* (Seo and Koshiba, 2002; Takino et al., 2019). The PYL1-ABI1 interaction is presumably specific to ABA and not triggered by plant intermediates. However, since PYL1 and ABI1 are likely not exposed to *B. cinerea* ABA intermediates in nature it is possible that these intermediates trigger the PYL1-ABI1 interaction. A previous study investigated if *B. cinerea* culture filtrate has an effect on the ABA-sensitive plant *Lemna minor* (Siewers, 2004). No effect was observed for the culture filtrate of a Δ*bcaba1 B. cinerea* strain, suggesting that ⍺-IE does not trigger the ABA response in plants. However, no other ABA pathway knockout strains were tested in the study and the effect of downstream intermediates remains elusive. In yMO28, the enzyme BcABA2 is missing from the ABA pathway, which led to the accumulation of a compound with m/z 235 in the supernatant, likely corresponding to the ABA intermediate α-IAA (Supp. Fig. S8). This compound was not detected in yMO27, lacking *bcaba1*. Both strains, yMO27 and yMO28, showed close to identical GFP outputs (Figure 5), leading to the conclusion that the m/z 235 compound does not trigger the ABA biosensor. The ligand specificity of the ABA biosensor could be further investigated by feeding ABA intermediates or by constructing a strain that lacks *bcaba4*, which should lead to accumulation of the intermediate DH-⍺-IAA.

As was expected, we observed a positive correlation between the GFP reporter signal and the RFP signal (Supp. Fig. S13), demonstrating that constitutive expression of *RFP* can be used to normalize the reporter output to extrinsic noise. The second fluorophore also enables more accurate distinction of sub-populations, assumed to be budded and un-budded cells, compared to using histograms (Figure 5A, Supp. Fig. S13).

In conclusion, we validated that the sensor response correlates with the ABA titers produced in yMO16 and yMO17 and demonstrated the usefulness of the p*THI4*-regulated design. For the ABA feeding experiment (Figure 3) and for this analysis (Figure 5), the timepoint of measurement has been identified as a crucial parameter when using the biosensor in future HTSs.

## Conclusion

Using high-throughput methods, millions of enzyme variants can be screened within hours and improved mutants can be isolated. Nonetheless, HTS approaches come with their own challenges, often suffering from a high percentage of false positives. In this study we aimed to address this issue by developing platform strains that are specifically engineered for this purpose. We developed screening platforms for two rate limiting enzymes in *S. cerevisiae* ABA cell factories, BcABA1 and BcABA2.

We thoroughly characterised and evaluated ABA biosensor variants and identified cultivation time as a critical parameter. Cultivation time did not only affect the biosensor output in ABA-producing strains (Figure 5), which was expected since ABA accumulates over time, but also had a considerable impact the biosensor response curve in cultures with exogenously added ABA (Figure 3). Cultivation time is arguably an often overlooked parameter in biosensor studies and single time point measurements are common. There are likely multiple reasons for the observed changes between the time points. Differences in metabolic activities between the glucose and ethanol phase likely contribute. Furthermore, variations in growth rate can play a substantial role, since they have global effects on cellular metabolism, not just on a biological level, e.g. due to changes in ribosome availability (Mayer and Grummt, 2006), but also on a physical level via the dilution of cellular components during cell growth and division (Hintsche and Klumpp, 2013). A recent study investigated the correlation between growth rate and dynamic range of biosensors in *E. coli* (Hartline and Zhang, 2022). The authors highlight that the correlation is biosensor-specific and is affected by factors unrelated to the sensor setup like transport of the ligand. To our knowledge, comparable studies in *S. cerevisiae* are missing so far, but would be especially valuable for industry-oriented applications of biosensors, e.g. for the dynamic regulation of metabolic pathways in cell factories (Hartline and Zhang, 2022; Tan and Prather, 2017). In recent years, there has been a push towards predictive modelling of biosensor characteristic in the synthetic biology community (Berset et al., 2017; Chen et al., 2018; Hartline and Zhang, 2022; Mannan et al., 2017; Shaw et al., 2019), which will help to further elucidate response curve dependencies that go beyond the genetic parts of the biosensor itself.

The constructed platform strains contain a simple genetic switch to regulate the expression level of *bcaba1* and *bcaba2* variants, thereby changing ABA production levels and biosensor output (Figure 1C, Figure 4, Figure 5). The ABA titer can be adjusted to the operational range of the biosensor and be optimized for library screenings. For optimal screening conditions, the thiamine concentration (and cultivation time) should be adjusted so that wild type enzymes show minimal reporter output, leaving enough response curve “headspace” to distinguish improved enzyme variants and reduce the number of false positives. Nonetheless, a thorough evaluation of the benefits of tuneable ABA production requires screening of a mutant library.

After a successful screening, the discovered enzyme variants can be further engineered, e.g. by combining beneficial mutations or by performing site saturation mutagenesis on amino acid residues that were shown to affect enzyme activity. The platform strains can be used to screen a second, more targeted library and higher thiamine concentrations can be used to compensate for the increased baseline activity of the enzymes. Tuneable ABA production could also prove useful for the in-depth characterization and comparison of different variants, potentially providing information about optimal expression levels and promising promoter candidates in ABA cell factories. In addition, the biosensors characterised in this study could be used in the ABA production strains, e.g. to separate growth and production phase in a bioreactor setup (Tan and Prather, 2017).

ABA titers in *S. cerevisiae* remain low in comparison to ABA-overproducing *B. cinerea* strains (Gong et al., 2014; Otto et al., 2019). Yet, the vast knowledge, engineering as well as screening techniques available for *S. cerevisiae* allows rapid prototyping and utilisation of biosensors and genetic circuits. Improved BcABA1 or BcABA2 variants, discovered using the described screening platforms, might either be directly transferable to *B. cinerea* or knowledge about mutation hotspots in the enzymes might indirectly benefit *B. cinerea* cell factories.

Our study represents an exploratory analysis of ABA biosensors in yeast and furthermore examines the concept and usefulness of screening platform strain.

## Acknowledgments

We would like to thank Lucy Fang-I Chao and Tyler Doughty for providing the p3036-pGAL1*-yeGFP expression cassette, as well as the gRNAs and repair fragments for the *GAL4* and *GAL80* deletions. Furthermore, we would like to acknowledge Louis Scott and Tim Eckerström for providing *THI4* promoter fragment. We would also like to thank Jing Fu and Michael Gossing for sharing their modified high efficiency transformation protocol with us.

## Contribution

MO, FD and VS conceived this project. MO, FD and VS designed the research. MO and YD performed and analysed the experiments. MO, YD, FD and VS prepared the manuscript.

