## Supplementary data for "Sense and Screen-ability: Development of tuneable, biosensor-based screening platforms for abscisic acid"

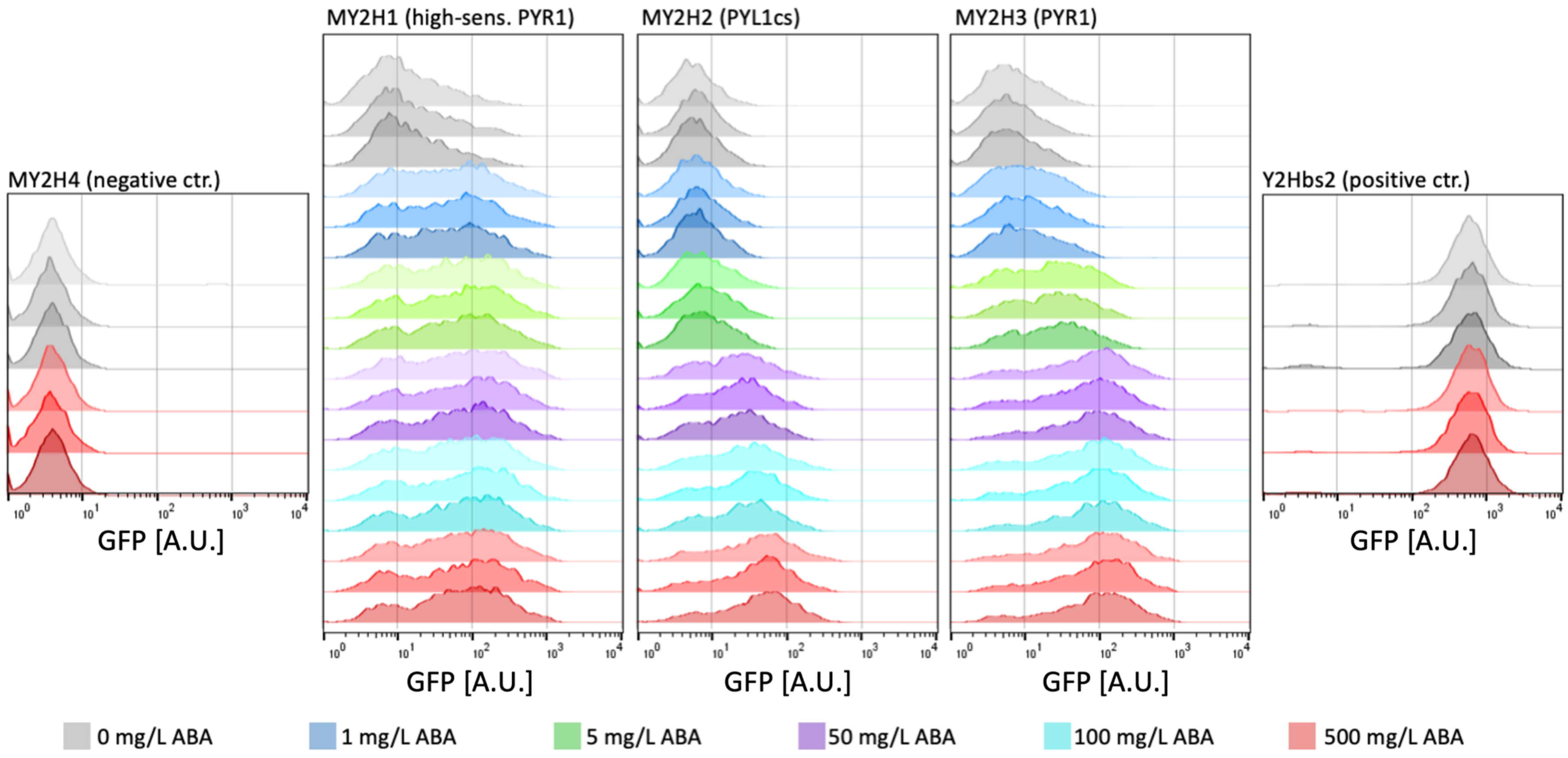


Figure S1: ABA biosensor responses in strains expressing the fusion proteins from a centromeric plasmid. The strains were grown in minimal media and ABA was added in varying concentrations (colours) at the beginning of the main culture. Flow cytometry was performed after 19 h. Three biological replicates are shown per strain and condition.

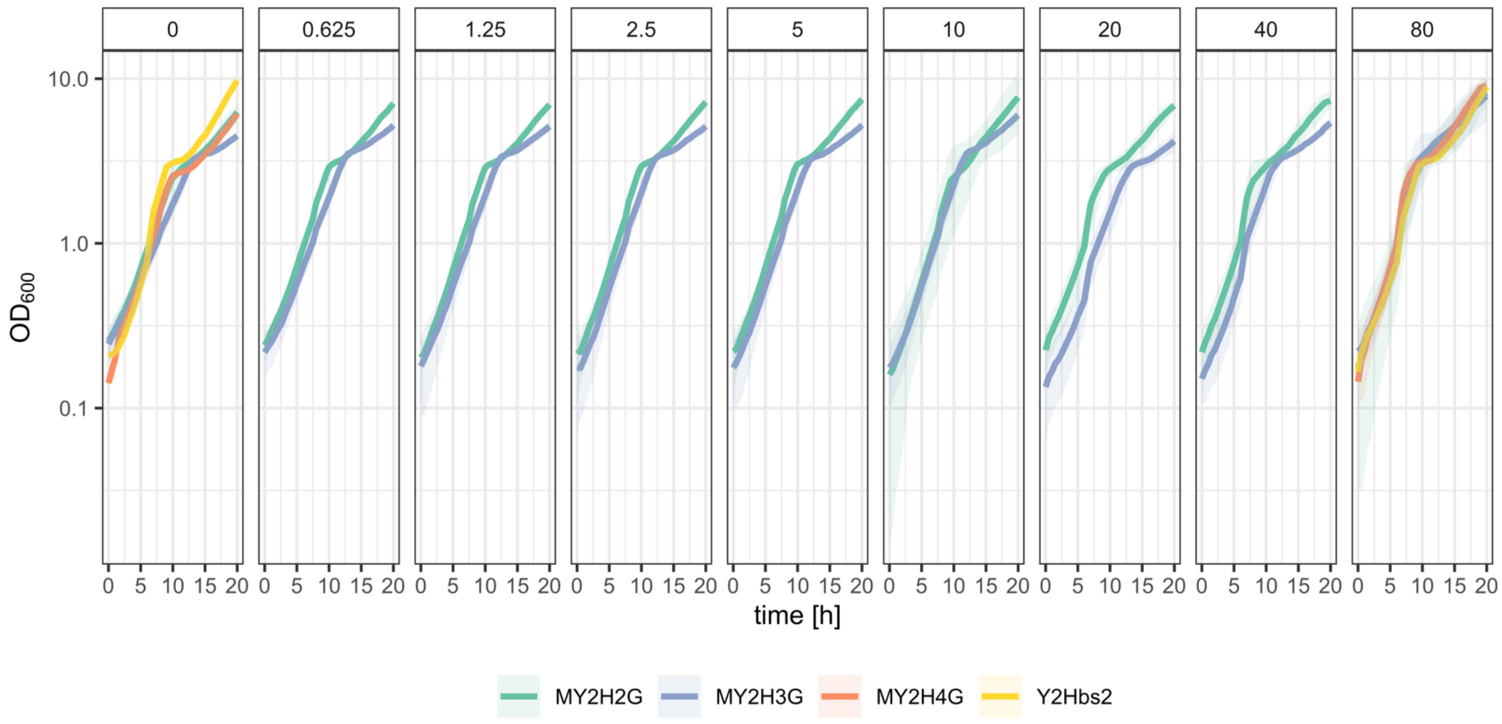


Figure S2: Growth profiles of biosensor strains, negative and positive control. A GrowthProfiler (Enzyscreen) was used to monitor the $OD_{600}$. Strains were grown in mineral media with varying concentrations of ABA (column labels in mg/L). Flow cytometry samples were taken after 6 h and 20 h (see Figure 3 in main document).

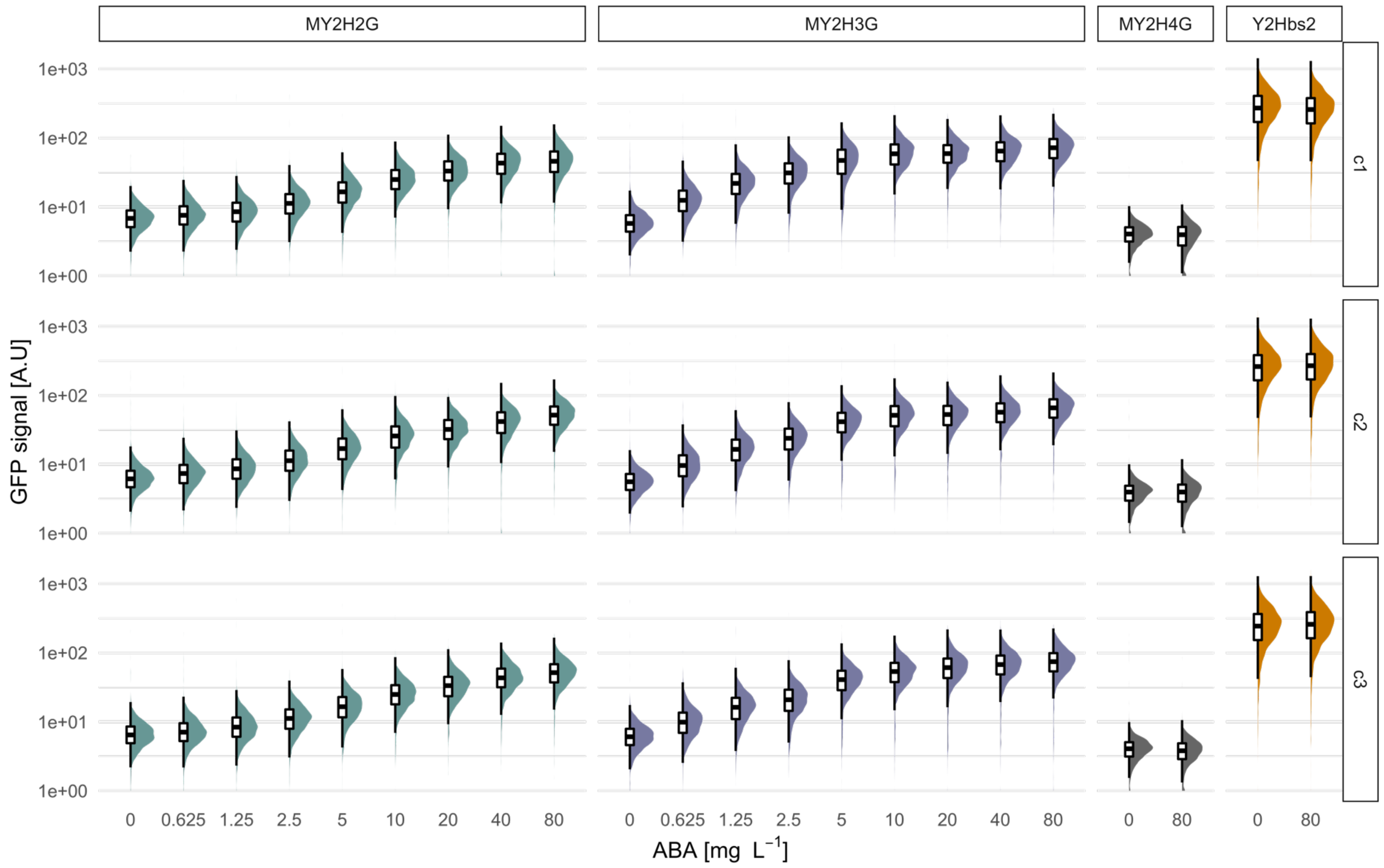


Figure S3: Flow cytometry analysis of the biosensor strains and control strains at 20 h. See Figure 3 in main text. Strains were grown in mineral media with varying concentrations of ABA. Biological replicates (clones c1 to c3) are shown in rows.

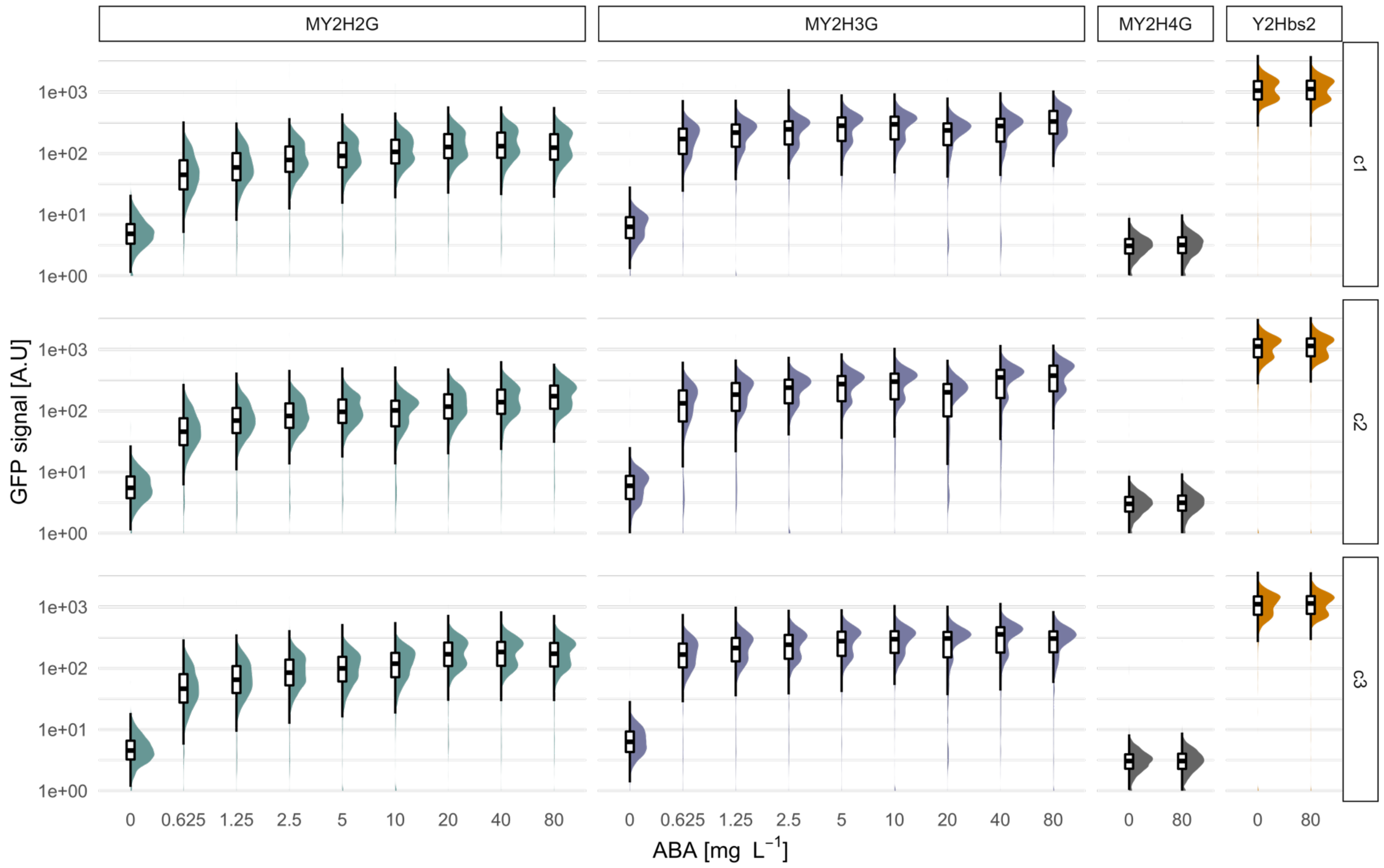


Figure S4: Flow cytometry analysis of the biosensor strains and control strains at 20 h. See Figure 3 in main text. Strains were grown in mineral media with varying concentrations of ABA. Biological replicates (clones c1 to c3) are shown in rows.

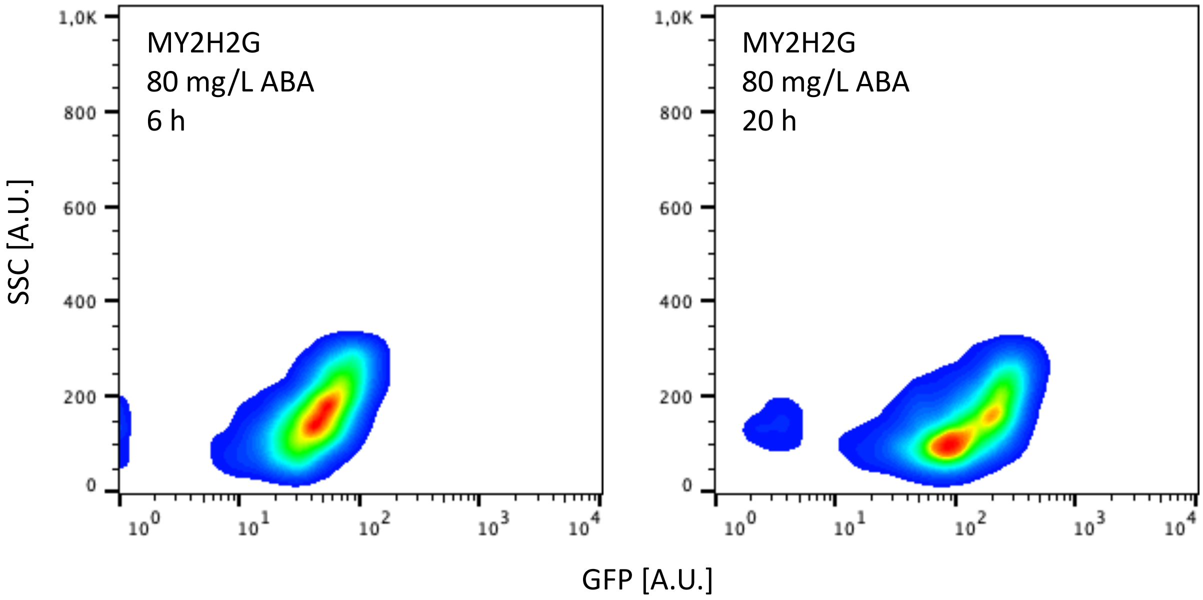


Figure S5: Correlation of SSC and GFP signal exemplified for MY2H2G induced with 80 mg/L and cultivated for 6 or 20 h (same experiment as in Figure S3 and S4). Two distinct populations are visible for the 20 h measurement and the GFP signal shows a positive correlation with the SSC signal. A similar GFP correlation was visible for FSC (data not shown).

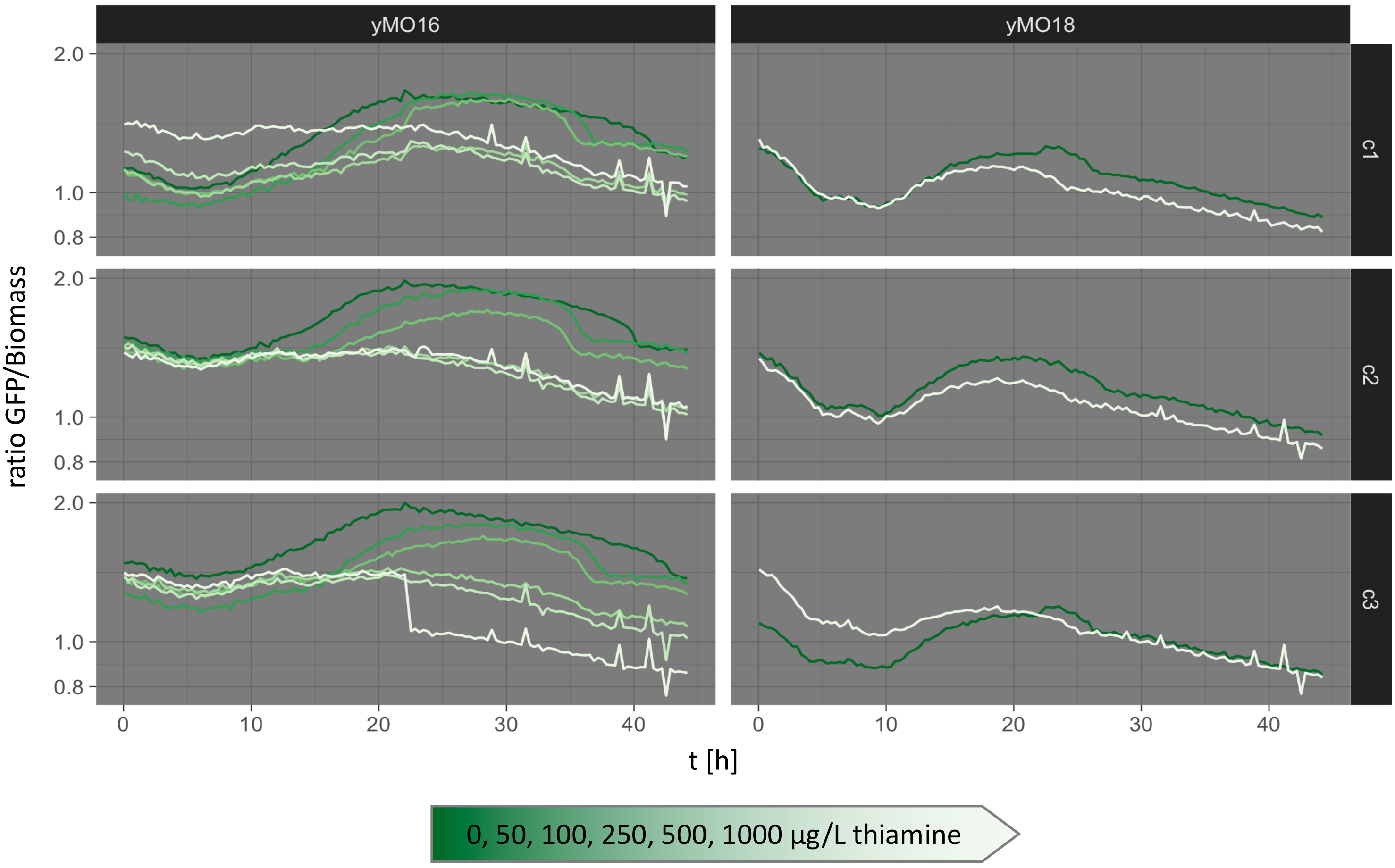


Figure S6: Ratio of GFP to Biomass detected over time in yMO16 and the negative control strain yMO18. A Biolector (M2P labs) was used to monitor fluorescence and growth. Cells were cultivated in SD media (unbuffered) with varying concentrations of thiamine (colour-coded, green = absence of thiamine, white = 2000 µg/L thiamine). The control strain yMO18 has the same genotype as yMO16 but carries an empty plasmid instead of pMMC33.

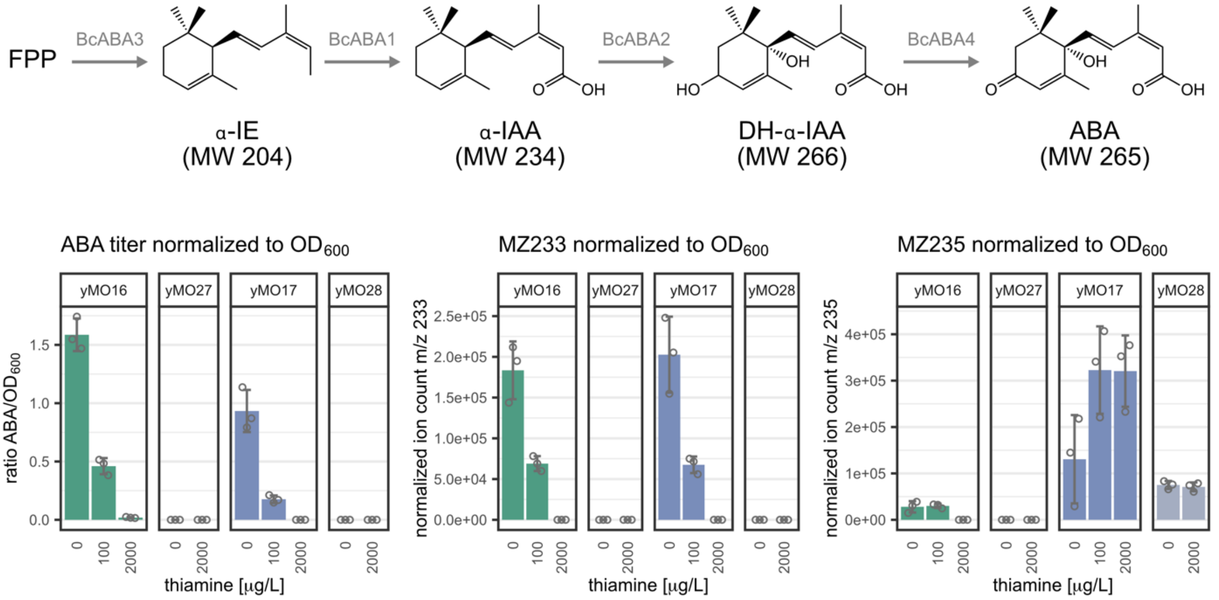


Figure S7: Normalized ABA titer and ion counts of potential ABA intermediates or side products with the mass-to-charge ratio (m/z) 233 or 235. The strains (colour-coded) were grown in various thiamine concentrations and HPLC analysis of the culture supernatant was performed after 20 h of cultivation. The mean and standard deviation was calculated from three biological replicates. Individual data points are shown as circles. The ABA pathway with molecular weights (MW) is shown above. Abbreviations: FPP = farnesyl pyrophosphate, α-IE = α-ionylideneethane, α-IAA = α-ionylideneacetic acid, DH-α-IE = 1′,4′-trans-dihydroxy-α-ionylideneacetic acid, ABA = abscisic acid

### Extended Results and Discussion: ABA pathway intermediates/side products:

MZ233 was previously identified as an ABA intermediate or side product in *S. cerevisiae* (Otto *et al.* 2019). No ABA or MZ233 was detected for the control strains yMO27 and yMO28. In yMO16 and yMO17 MZ233 titers roughly correlate with ABA production. We previously presumed that MZ233 originates downstream of BcABA1 and upstream of BcABA2, however, this does not seem to be the case since BcABA2 appears to be required for MZ233 formation. BcABA2 catalyses two oxidation steps and ABA side products have been reported in *Aspergillus oryzae*, where they seem to originate from partially oxidized α-IAA (Takino *et al.* 2019). This could also be the case in *S. cerevisiae*. Alternatively, MZ233 could be a degradation product of ABA or DH-α-IAA.

MZ235 accumulated in yMO17 and yMO28, strains in which the BcABA2 reaction step is limiting or lacking. This is in line with the hypothesis that the detected ion is the ABA intermediate α-IAA. Small amounts of MZ235 were detectable in yMO16 when cultivated in the absence of thiamine or in 100 µg/L thiamine. No MZ235 was detected for yMO16 in 2000 µg/L thiamine when p*THI4-bcaba1* is repressed, confirming that it is a downstream intermediate.

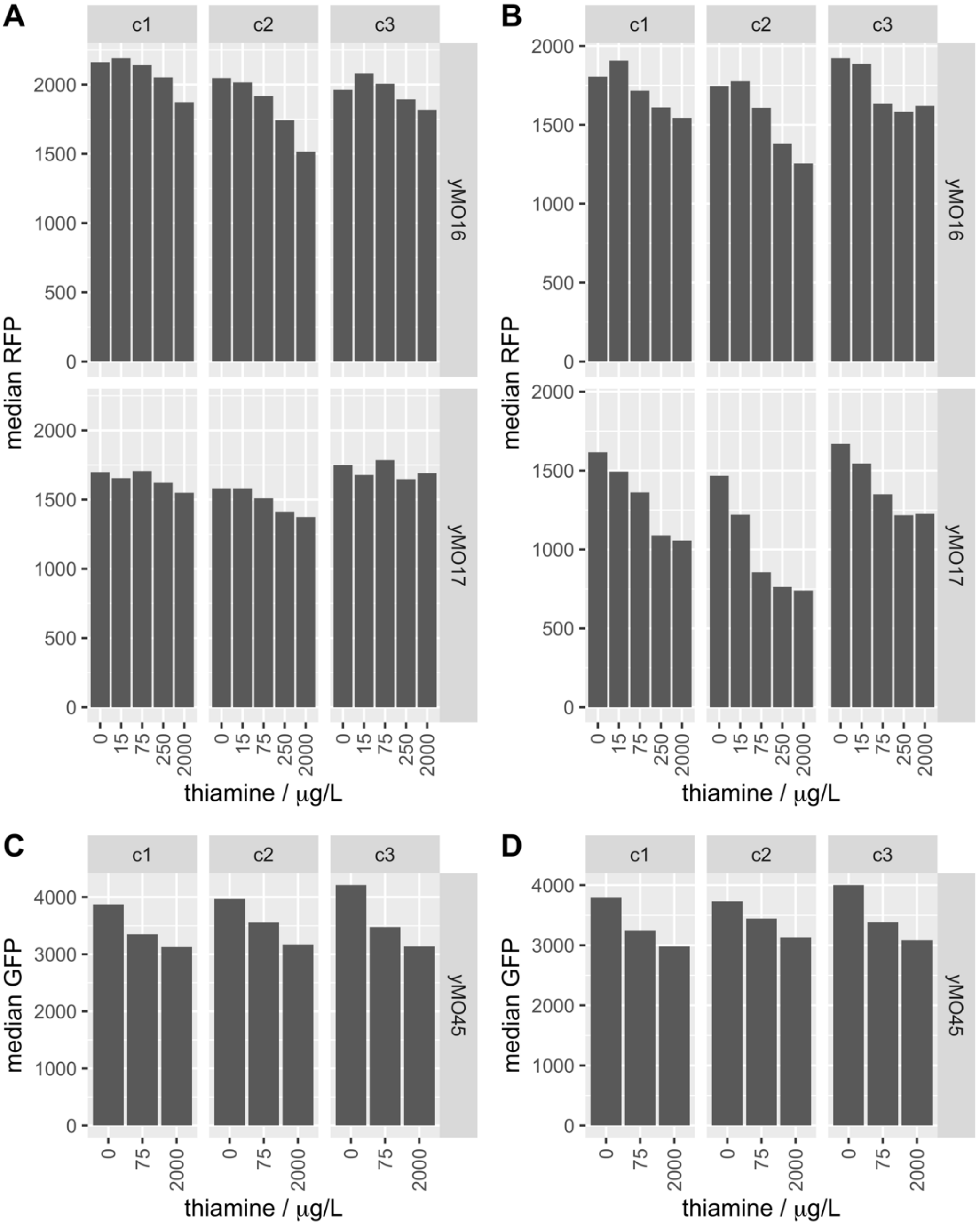


Figure S8: Median fluorescence of constitutively expressed fluorescent proteins dependant on thiamine concentration. Three biological replicates (clones c1 to c3) are shown. See Figures S8-S11 for the complete data sets. A: Median RFP signal of the strains yMO16 and yMO17 after 19 h of cultivation. B: Median RFP signal of the strains yMO16 and yMO17 after 23 h of cultivation. C: Median GFP signal of a control strain yMO45 (strain Y2Hbs2 with empty plasmid) with constitutive expression of p*GAL1*-yeGFP* after 19 h of cultivation. D: Median GFP signal of a control strain yMO45 (strain Y2Hbs2 with empty plasmid) with constitutive expression of p*GAL1*-yeGFP* after 23 h of cultivation.

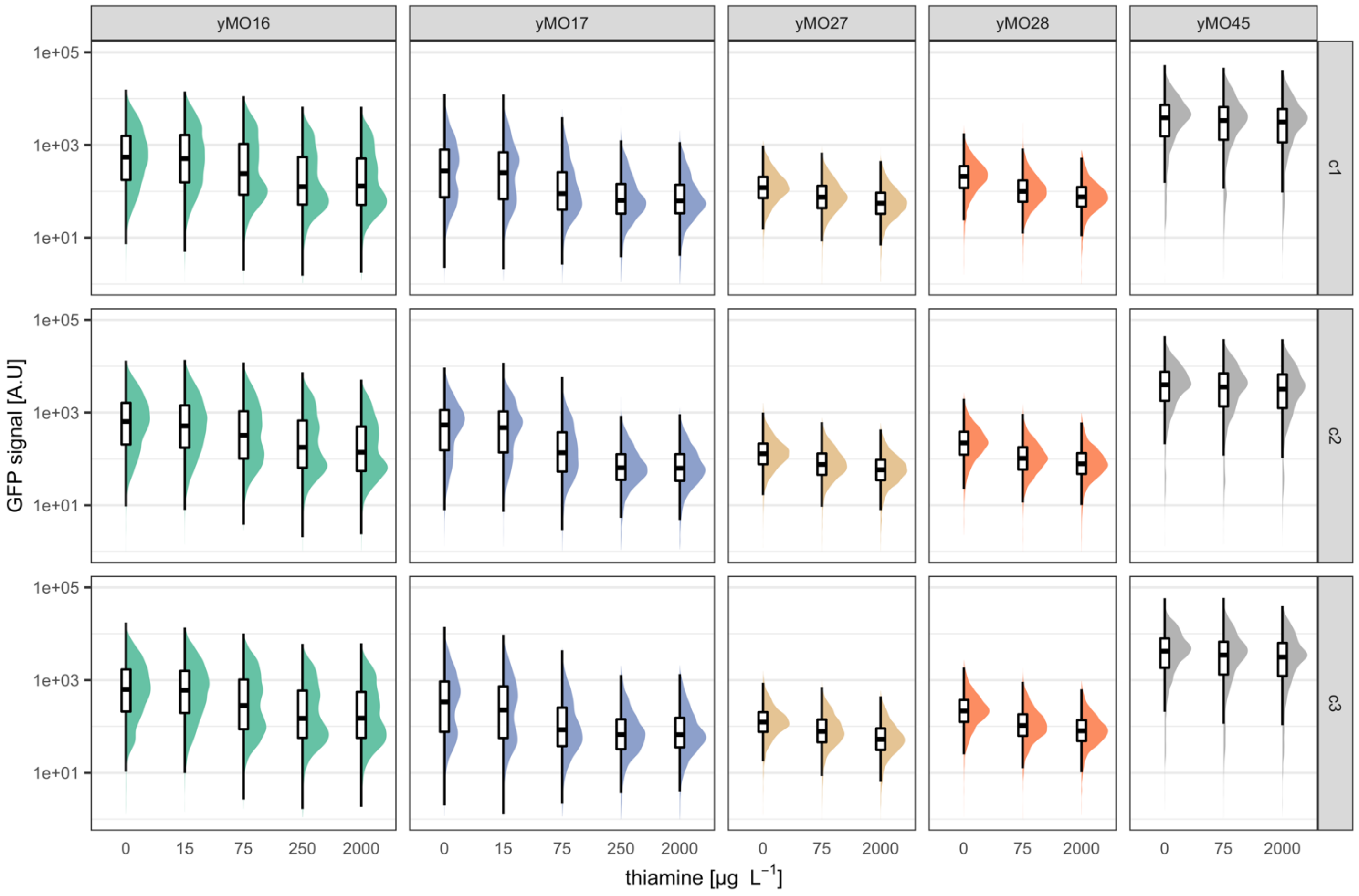


Figure S9: GFP reporter output for yMO16, yMO17, yMO27, yMO28 and yMO45 after 19 h of cultivation. The control strain yMO45 (strain Y2Hbs2 with empty plasmid) expresses the native *GAL4* gene leading to constitutive expression of p*GAL1**-yeGFP*. Strains were grown in buffered SD media (pH 6) with varying concentrations of thiamine. Three biological replicates (clones c1 to c3) are shown in rows.

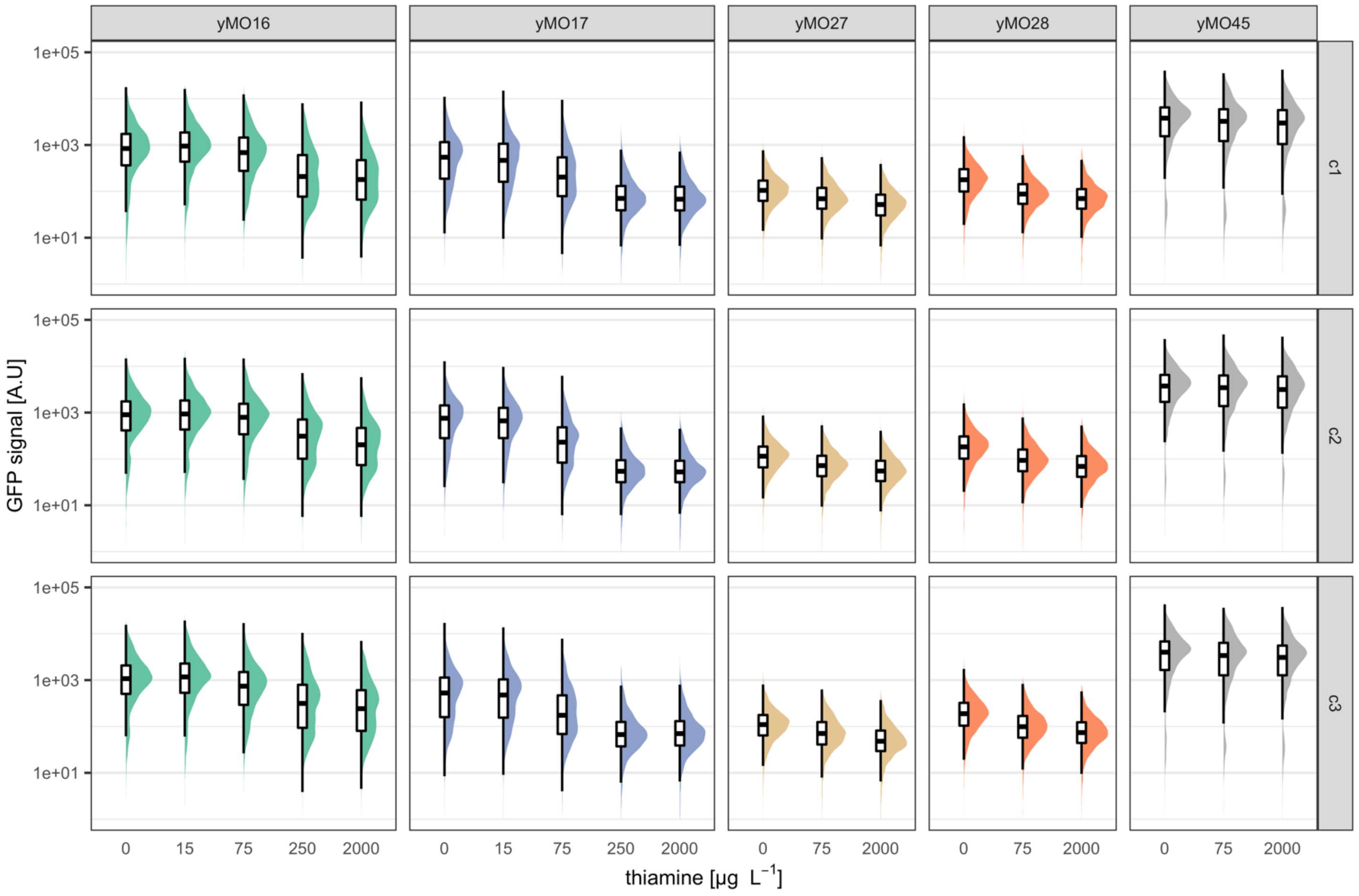


Figure S10: GFP reporter output for yMO16, yMO17, yMO27, yMO28 and yMO45 after 23 h of cultivation. The control strain yMO45 (strain Y2Hbs2 with empty plasmid) expresses the native *GAL4* gene leading to constitutive expression of p*GAL1*-yeGFP*. Strains were grown in buffered SD media (pH 6) with varying concentrations of thiamine. Three biological replicates (clones c1 to c3) are shown in rows.

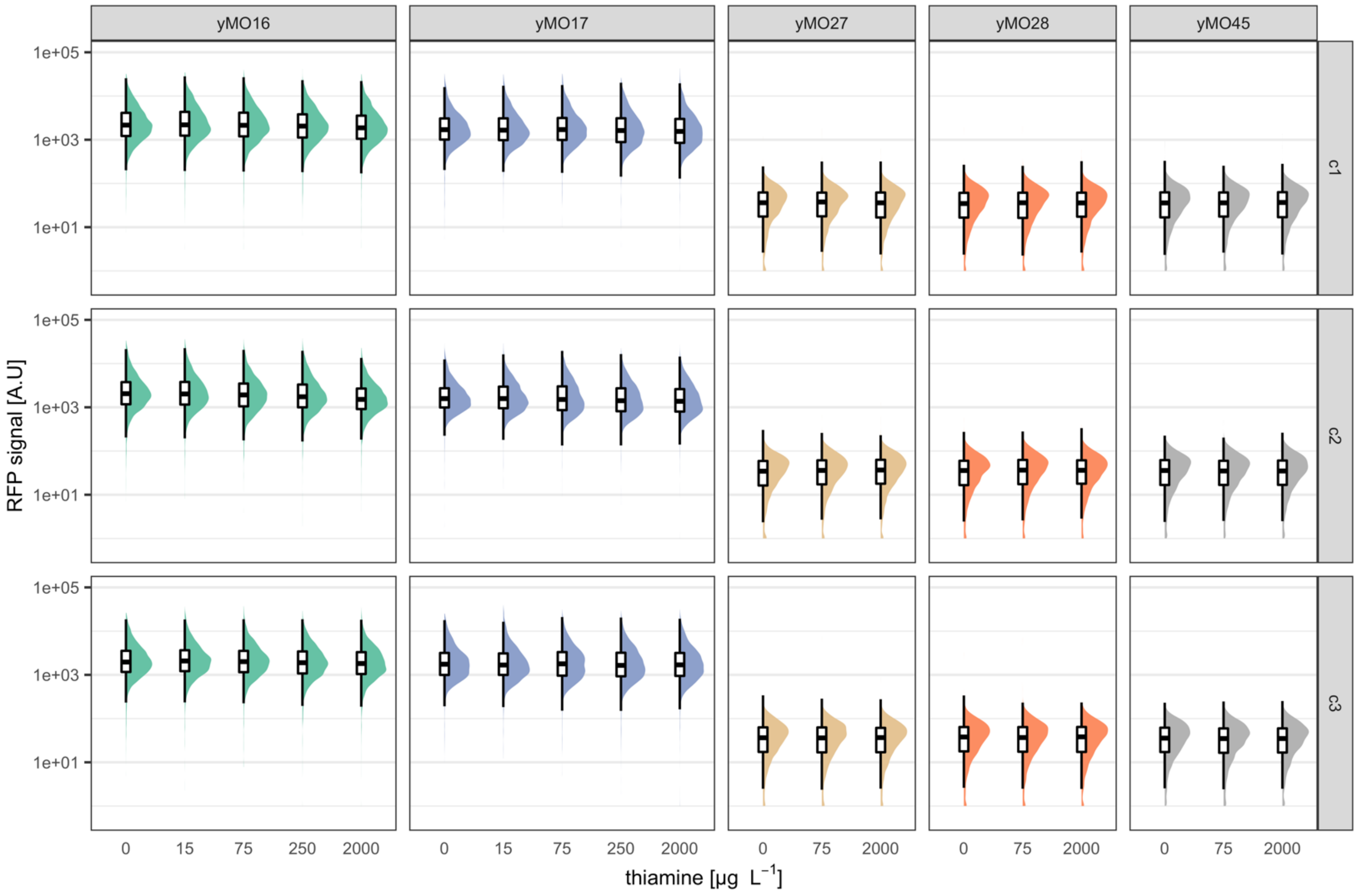


Figure S11: RFP signal of yMO16, yMO17, yMO27, yMO28 and yMO45 after 19 h of cultivation. The ABA-producing strains yMO16 and yMO17 constitutively express *miRFP670*. The strains yMO27, yMO28 and yMO45 (strain Y2Hbs2 carrying an empty plasmid) do not contain the *miRFP670* gene. Strains were grown in buffered SD media (pH 6) with varying concentrations of thiamine. Three biological replicates (clones c1 to c3) are shown in rows.

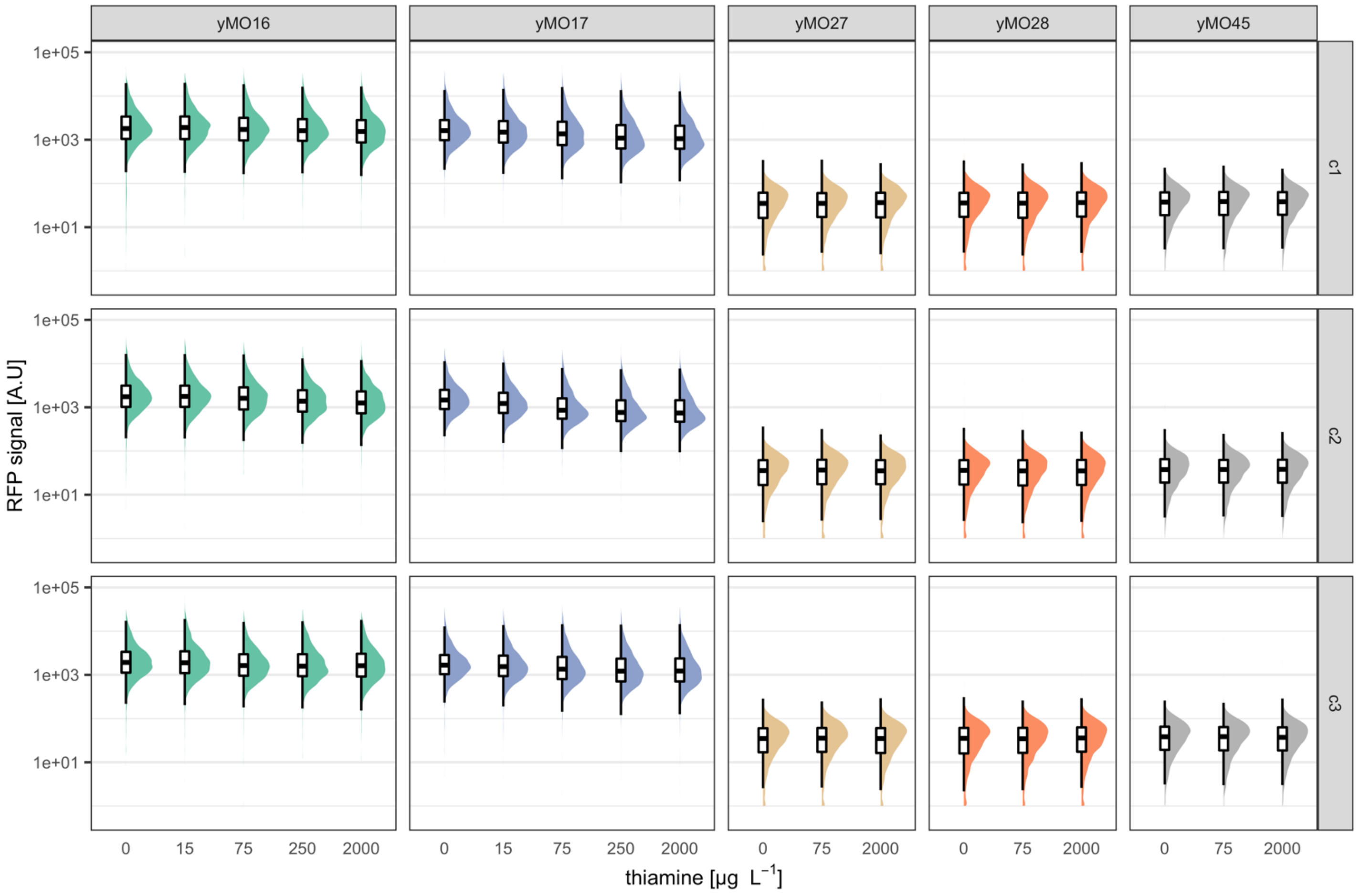


Figure S12: RFP signal of yMO16, yMO17, yMO27, yMO28 and yMO45 after 23 h of cultivation. The ABA-producing strains yMO16 and yMO17 constitutively express *miRFP670*. The strains yMO27, yMO28 and yMO45 (strain Y2Hbs2 carrying an empty plasmid) do not contain the *miRFP670* gene. Strains were grown in buffered SD media (pH 6) with varying concentrations of thiamine. Three biological replicates (clones c1 to c3) are shown in rows.

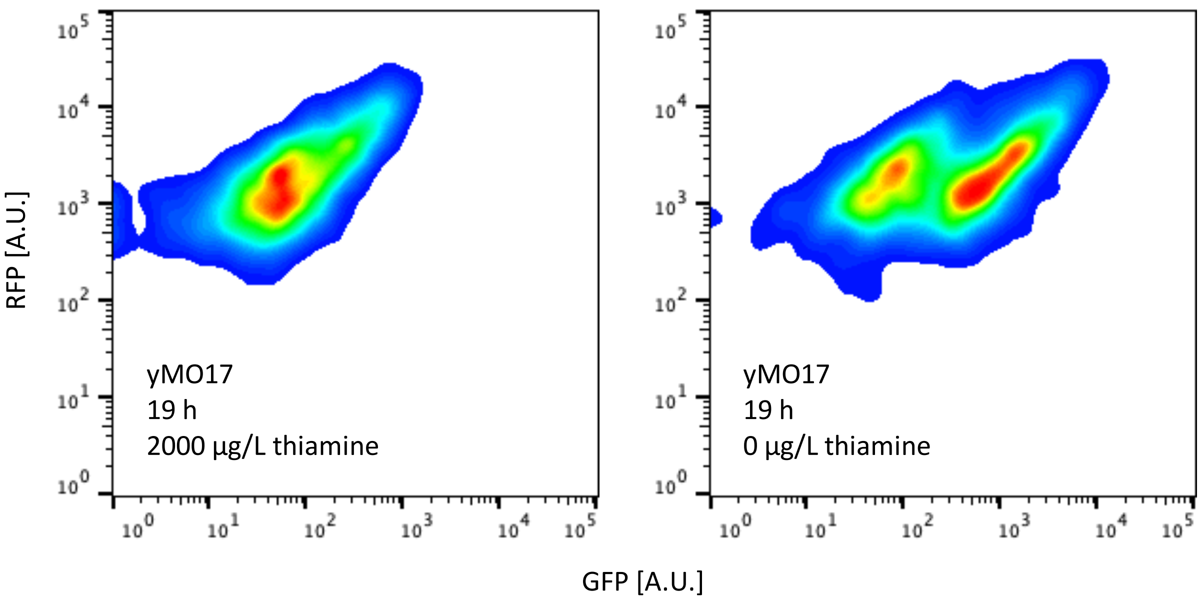


Figure S13: Correlation of RFP and GFP signal exemplified for yMO17 grown in absence or 2000 µg/L thiamine for 19 h (same experiment as in Figure S9-S11). GFP and RFP have a positive correlation indicating that RFP can be used to normalize the biosensor output for extrinsic noise.

Table S1: Primers

| Primer name | Sequence |
|---|---|
| 175 | TACCAAACGACGAGCGTGACACCACGATG |
| 176 | GTCACGCTCGTCGTTTGGTATGGCTTCATTCAGC |
| 178 | AATGCGTGCGGCCGGCCGCACACACCAT |
| 179 | TGCGGCCGGCCGCACGCATTCCATGCGAG |
| 185 | CCTGCAGCCCGGGGGATCTTATCTTGGAACTTCTTTTAACATAAC |
| 196 | GCAAAGCTCGCCAGTGCTCCA |
| 198 | AGGCCGGCCGCACACACC |
| 237 | GCATCGTCTCATCGGTCTCATATGGATAAAGCGGAATTAATTCCCG |
| 238 | ATGCCGTCTCAGGTCTCAAGAACCCTCTTTTTTTGGGTTTGGTG |
| 239 | GCATCGTCTCATCGGTCTCATATGAAGCTACTGTCTTCTATCG |
| 240 | ATGCCGTCTCAGGTCTCAAGAACCCGATACAGTCAACTGTCTTTG |
| 364 | GCATCGTCTCATCGGTCTCATATGGCTGCAGACCAATTGGT |
| 365 | ATGCCGTCTCAGGTCTCAGGATCCTTAGGATTTAATGCAGGTGAC |

Table S2: PCR reactions

| **Primer pair** | **Template** | **Polymerase** | **$T_{ann}$** | **$t_{elo}$ (min:sec)** | **Use** |
|---|---|---|---|---|---|
| 175/178 | pCfB2909-ABA2-ABA4 (Otto *et al.* 2019) | Phusion | 72° | 3:30 | Gibson assembly of pCfB2909-ABA4 |
| 176/179 | pCfB2909-ABA2-ABA4 (Otto *et al.* 2019) | Phusion | 71° | 1:30 | Gibson assembly pCfB2909-ABA4 |
| 176/197 | pCfB2904-ABA1-CPR (Otto *et al.* 2019) | Phusion | 64° | 1:30 | Gibson assembly pCfB2903-CPR |
| 185/198 | pCfB2904-ABA1-CPR (Otto *et al.* 2019) | Phusion | 72° | 5:00 | Gibson assembly pCfB2903-CPR |
| 237/238 | pGBDU-C1 (James, Halladay and Craig 1996) | Phusion | 53° | 0:30 | amplify Gal4BD |
| 239/240 | pGBAU-C1 (James, Halladay and Craig 1996) | Phusion | 53° | 0:30 | amplify Gal4AD |
| 364/365 | XI5b-tHMG1 (López *et al.* 2015) | Primestar | 55° | 2:15 | amplify tHMG1 |
| 397/398 | pMMC33 | PrimeStar | 53° | 7:30 | amplify plasmid backbone for *bcaba1* library |
| 398/400 | pMMC34 | PrimeStar | 61° | 7:30 | amplify plasmid backbone for *bcaba2* library |

Table S3: List of MoClo assemblies (Lee et al. 2015; Otto et al. 2021)

| Plasmids name | MoClo parts used | MoClo level | Details |
|---|---|---|---|
| pMC3a-Gal4AD | pYTK001, PCR 237/238 | level-0 | |
| pMC3a-Gal4BD | pYTK001, PCR 239/300 | level-0 | |
| pMC3b-PYLcs | pYTK001, gene fragment | level-0 | genes fragments were ordered from Genscript |
| pMC3b-PYR1 | pYTK001, gene fragment | level-0 | genes fragments were ordered from Genscript |
| pMC3b-PYR1 $^{F61L\_A160C}$ | pYTK001, gene fragment | level-0 | genes fragments were ordered from Genscript |
| pMC3b-ABIcs $^{D413L}$ | pYTK001, gene fragment | level-0 | genes fragments were ordered from Genscript |
| pMC3b-ABI1 $^{D413L}$ | pYTK001, gene fragment | level-0 | genes fragments were ordered from Genscript |
| pMC3-tHMG1 | pYTK001, PCR 364/365 | level-0 | |
| pXI5-MMC5K | pMC-XI5, pYTK012, pYTK055, pMC3a-Gal4BD, pMC3b-ABI1 $^{D413L}$ | level-1 | |
| pMMC5 | pMC-Ura-Cen, pYTK012, pYTK055, pMC3a-Gal4BD, pMC3b-ABI1 $^{D413L}$ | level-1 | |
| pMMC6 | pMC-Ura-Cen, pYTK012, pYTK055, pMC3a-Gal4BD, pMC3b-ABI1cs $^{D413L}$ | level-1 | |
| pMMC7 | pMC-His-Cen, pYTK013, pYTK053, pMC3a-Gal4AD, pMC3b-PYR1 | level-1 | |
| pMMC8 | pMC-His-Cen, pYTK013, pYTK053, pMC3a-Gal4AD, pMC3b- PYR1 $^{F61L\_A160C}$ | level-1 | |
| pMMC9 | pMC-His-Cen, pYTK013, pYTK053, pMC3a-Gal4AD, pMC3b-PYLcs | level-1 | |
| pMMC16 | pYTK009, pYTK056, pMC3-bcaba1, pMC-Ura-Cen | level-1 | |
| pMMC19 | pMC3-pTHI4, pYTK055, pMC3-bcaba2, pMC-His-Cen | level-1 | |
| pMMC17 | pYTK010, pYTK055, pMC3-bcaba2, pMC-His-Cen | level-1 | |
| pMMC18 | pMC2-pTHI4, pYTK056, pMC3-bcaba1, pMC-Ura-Cen | level-1 | |
| pMMC22 | pMC-Ura-Cen, pYTK009, pYTK054, pMC3-tHMG1 | level-1 | |
| pMMC32 | pMC-His-Cen, pYTK013, pYTK055, pMC3-miRFP670 | level-1 | |
| pXI5-MMC12K | pMC-XI5, pMMC5, pMMC8 | level-2 | |
| pXI5-MMC11K | pMC-XI5, pMMC5, pMMC7 | level-2 | |
| pXI5-MMC10K | pMC-XI5, pMMC6, pMMC9 | level-2 | |
| pMMC33 | pMMC17, yMO18, pMC-Ura-Cen-multi | level-2 | |
| pMMC34 | pMMC16, yMO19, pMC-Ura-Cen-multi | level-2 | |
| pX3-tHMG1-miRFP | pMC-X3, pMMC22, pMMC32 | level-2 | |
| pMC-Ura-Cen-multi | pYTK008, pYTK047, pYTK073, pYTK074, pYTK081, pYTK084 | level-2 backbone | contains GFP dropout cassette |

Table S4: Media compositions

| | |
|---|---|
| **YPD** | |
| yeast extract (Merck) | 10 g/L |
| peptone from meat (Merck) | 20 g/L |
| glucose (Merck) | 20 g/L |
| for plates: agar agar (Merck) | 20 g/L |
| **SD (pH adjusted to 6 with KOH)** | |
| complete supplement mix dropout uracil (Formedium) | 0.77 g/L |
| yeast-nitrogen base without amino acids and thiamine (Formedium) | 6.9 g/L |
| glucose (Merck) | 20 g/L |
| for plates: agar agar (Merck) | 20 g/L |
| for buffered liquid media: 1 M $Na_2HPO_4$ solution | 63.1 mL/L |
| for buffered liquid media: 1 M citric acid solution | 18.45 mL/L |
| **LB (pH adjusted to 6.5 with NaOH)** | |
| peptone from casein (Merck) | 10 g/L |
| NaCl (Merck) | 10 g/L |
| yeast extract (Merck) | 5 g/L |
| for plates: agar agar (Merck) | 20 g/L |
| **Mineral media (pH adjusted to 6.5 with KOH)** | |
| ammonium sulfate (Merck) | 7.5 g/L |
| monopotassium phsophate (Merck) | 14.4 g/L |
| magnesium sulfate heptahydrate (Merck) | 0.5 g/L |
| glucose (Merck) | 20 g/L |
| trace metal solution | 2 mL/L |
| vitamin solution | 1 mL/L |
| for plates: agar agar (Merck) | 20 g/L |
| uracil (Alfa Aesar) | 100 mg/L |
| histidine (Alfa Aesar) | 120 mg/L |
| **Trace metal solution** | |
| $FeSO_4 \cdot 7H_2O$ | 3 g/L |
| $ZnSO_4 \cdot 7H_2O$ | 4.5 g/L |
| $CaCl_2 \cdot 2H_2O$ | 4.5 g/L |
| $MnCl_2 \cdot 4H_2O$ | 1 g/L |
| $CoCl_2 \cdot 6H_2O$ | 300 mg/L |
| $CuSO_4 \cdot 5H_2O$ | 300 mg/L |
| $Na_2MoO_4 \cdot 2H_2O$ | 400 mg/L |
| $H_3BO_3$ | 1 g/L |
| KI | 100 mg/L |
| $Na_2EDTA \cdot 2H_20$ | 19 g/L |
| **Vitamin solution** | |
| D-Biotin | 50 mg/L |
| D-Pantothenic acid hemicalcium salt | 1 g/L |
| thiamine-HCl | 1 g/L |
| nicotinic acid | 1 g/L |
| pyridoxin-HCl | 1 g/L |
| 4-aminobenzoic acid | 0.2 g/L |
| myo-Inositol | 25 g/L |

Table S5: DNA oligos for gene knockouts

| **Description** | **Sequence** |
|---|---|
| Δ*gal80* gRNA fwd | TGCGCATGTTTCGGCGTTCGAAACTTCTCCGCAGTGAAAGATAAATGATCGTTTATAAAAGTAACATGATGTTTTAGAGCTAGAAATAGCAAGTTAAAATAAGGCTAGTCCGTTATCAAC |
| Δ*gal80* gRNA rev | GTTGATAACGGACTAGCCTTATTTTAACTTGCTATTTCTAGCTCTAAAACATCATGTTACTTTTATAAACGATCATTTATCTTTCACTGCGGAGAAGTTTCGAACGCCGAAACATGCGCA |
| Δ*gal80* repair fwd | TCCTTGCCGACCAGCGTATACAATCTCGATAGTTGGTTTCCCGTTCTTTCCACTCCCGTCAAGCATCTTGCCCTGTGCTTGGCCCCCAGTGCAGCGAACGTTATAAAAACGAATACTGAG |
| Δ*gal80* repair rev | CTCAGTATTCGTTTTTATAACGTTCGCTGCACTGGGGGCCAAGCACAGGGCAAGATGCTTGACGGGAGTGGAAAGAACGGGAAACCAACTATCGAGATTGTATACGCTGGTCGGCAAGGA |
| Δ*gal4* gRNA fwd | TGCGCATGTTTCGGCGTTCGAAACTTCTCCGCAGTGAAAGATAAATGATCAACAATTCCAGGCAAAATATGTTTTAGAGCTAGAAATAGCAAGTTAAAATAAGGCTAGTCCGTTATCAAC |
| Δ*gal4* gRNA rev | GTTGATAACGGACTAGCCTTATTTTAACTTGCTATTTCTAGCTCTAAAACATATTTTGCCTGGAATTGTTGATCATTTATCTTTCACTGCGGAGAAGTTTCGAACGCCGAAACATGCGCA |
| Δ*gal4* repair fwd | GTGTCTACGTAATGCACGCCATCATTTTAAGAGAGGACAGAGAAGCAAGCCTCCTGAAAGAATGAATCGTAGATACTGAAAAACCCCGCAAGTTCACTTCAACTGTGCATCGTGCACCAT |
| Δ*gal4* repair rev | ATGGTGCACGATGCACAGTTGAAGTGAACTTGCGGGGTTTTTCAGTATCTACGATTCATTCTTTCAGGAGGCTTGCTTCTCTGTCCTCTCTTAAAATGATGGCGTGCATTACGTAGACAC |
